# How Awakenings Shape Dream Recall: A Multilevel Study

**DOI:** 10.64898/2026.05.20.722678

**Authors:** Somayeh Ataei, Mahdad Jafarzadeh Esfahani, Nikolai Axmacher, Martin Dresler, Sarah F. Schoch

## Abstract

Dream recall varies substantially both between individuals and from night to night within the same individual. Although nocturnal awakenings are thought to facilitate the encoding and later retrieval of dream experiences, it remains unclear whether dream recall is shaped primarily by awakening frequency or by more specific awakening characteristics, including duration, sleep stage, and timing within the night. Here, we analyzed two cohorts: cohort 1 consisted of 708 adults spanning the full range of dream recall frequency, assessed across three waves with home sleep recordings and questionnaire-based dream recall frequency measures; cohort 2 consisted of 124 adults with high dream recall frequency, assessed across multiple nights with home sleep recordings and daily dream reports. Using multilevel models with within-between decomposition, we examined trait-like and state-like associations between awakening measures and dream recall outcomes.

At the trait level, both questionnaire-based dream recall frequency in cohort 1 and daily dream recall (i.e., a sense of having dreamed) in cohort 2 were associated with a specific nocturnal awakening profile: more habitual long REM awakenings and short NREM awakenings, with REM awakening effects remaining robust after adjustment for sleep duration. At the state level, in cohort 2, nights with more short and medium REM awakenings than usual increased the likelihood of morning dream recall, whereas nights with more long REM awakenings than usual increased the likelihood of morning dream content recall (i.e., remembering dream content). These findings support the arousal-retrieval and functional state-shift models, while highlighting important nuances in the associations between nocturnal awakenings and different dream recall outcomes.

## Introduction

The ability to remember dreams varies remarkably both between and within individuals. Given that humans likely dream every night, classical models of dream recall suggest that this variability is not due to differences in dream generation, but rather to factors that influence the encoding and subsequent retrieval of dream experiences (Fig. 1; for a review, see [1]). While some models emphasize dream features (the salience model [2] and the repression model [3]) or individual traits of the dreamer (the lifestyle model [4]) as factors influencing the encoding and retrieval of dreams, other models focus directly on encoding and retrieval processes, emphasizing neurophysiological and cognitive mechanisms occurring during and immediately after awakening from a dream (the arousal-retrieval model [5], the functional state-shift model [6], and the interference model [7]).

**Fig. 1:**
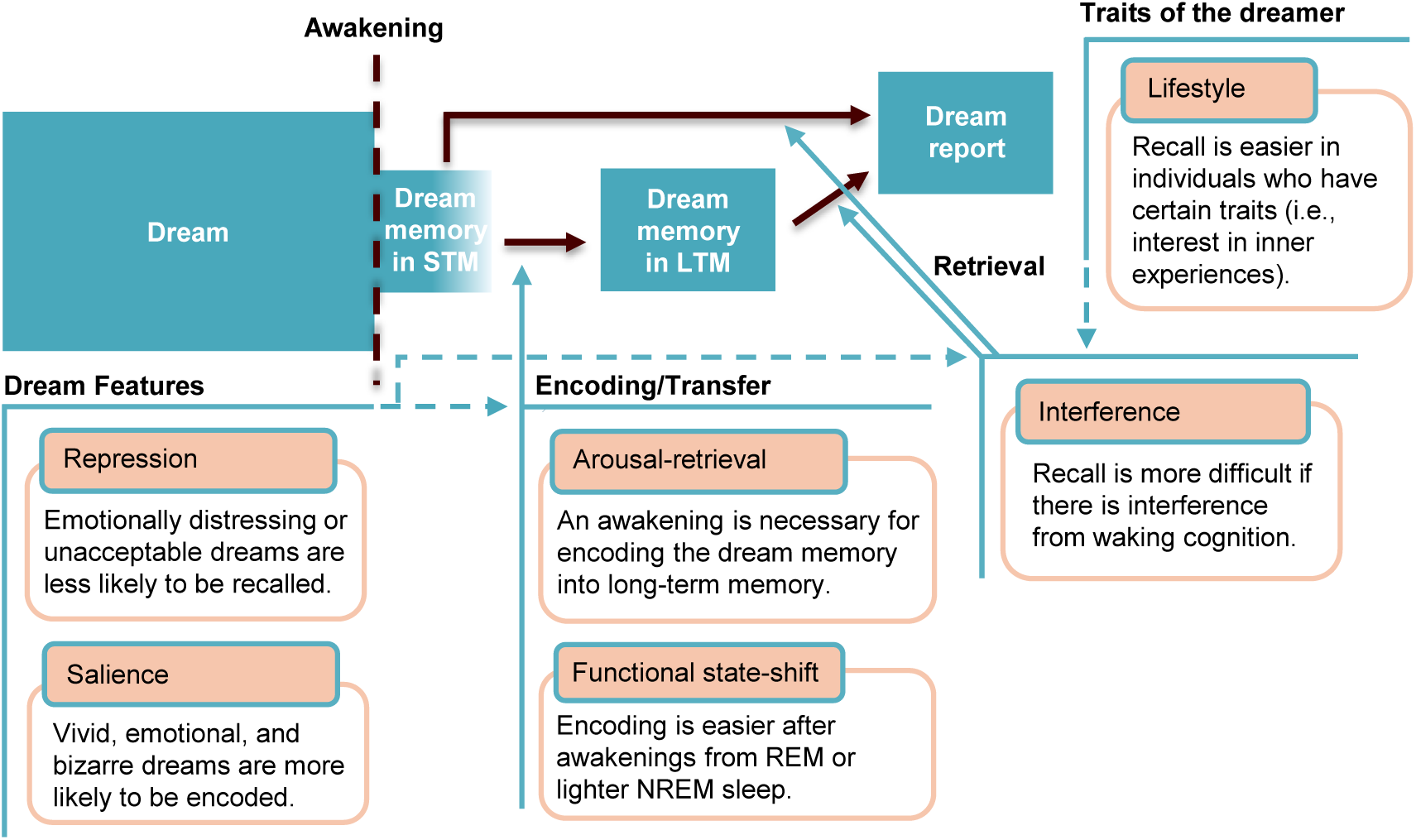
Dream recall process and models Schematic model of the dream recall process and major dream recall models. Encoding refers to the transfer of dream experiences from short-term memory (STM) to long-term memory (LTM), which occurs only if an awakening happens from the dream, according to the arousal-retrieval model [5]. Dream recall models can be grouped by the factors they emphasize: dream features (the salience model [2] and the repression model [3]); encoding (the arousal-retrieval model [5] and the functional state-shift model [6]), retrieval (the interference model [7]); and individual traits of the dreamer (the lifestyle model [4]). Blue dashed arrows highlight the interactions among these factors.

Despite theoretical differences, many models converge on the importance of nocturnal awakenings for the encoding and retrieval of dream experiences. The arousal–retrieval model posits that awakening provides the level of arousal required to transfer available dream material from short-term to long-term memory [5]. It has been suggested that this elevated arousal facilitates the attentional processes required for the retention of dream experiences [8] and that awakening reverses the aminergic demodulation and deactivation of the prefrontal cortex during sleep, thereby enabling effective retention of dream experiences [9]. Moreover, research on learning during sleep has shown that stimuli presented during sleep are recalled only if an awakening occurs within seconds of presentation (for a review, see [10]).

Empirical evidence broadly supports the importance of nocturnal awakenings for dream recall. Studies that combined one-night polysomnography (PSG) recordings—to objectively measure awakening frequency—with retrospective measures (e.g., dream questionnaires) have shown that individuals with high dream recall frequency experience more frequent awakenings than individuals with low dream recall frequency [11–13]. Dream recall is also significantly reduced following a night of total sleep deprivation, potentially because subsequent recovery sleep may contain fewer awakenings, and thus fewer opportunities for dream encoding [14]. Subjective measures of awakening frequency, which reflect awakenings that individuals notice and remember, have also been examined in a few studies [15–17]. In one of these, individuals with insomnia, who by definition report frequent nocturnal awakenings, were shown to report higher dream recall frequency (DRF) [15]. Two other studies showed that self-reported awakening frequency is positively correlated with DRF in healthy individuals [16,17].

Beyond how often awakenings occur, a key unresolved question is which *characteristics* of awakenings most strongly determine whether dreams are remembered. The only two studies examining this reported that individuals with high dream recall frequency experience more *long* awakenings (≥1 minute [12] or ≥2 minutes [13]) than individuals with low dream recall frequency. One of these studies further suggested that the minimum duration for an awakening to allow for memory encoding is approximately 2 minutes [12]. However, this inference was based on a comparison of average awakening durations between high- and low-frequency dream recaller groups (approximately 2 minutes vs. 1 minute). The second study focused exclusively on the frequency of longer awakenings (≥2 minutes), without considering shorter awakenings [13].

Moreover, although neither the arousal–retrieval model [5] nor the interference model [7] specifies an optimal awakening duration for dream recall, their integration suggests that recall may be favored by awakenings of intermediate duration—long enough to permit encoding, yet short enough to limit interference from waking cognition. While the first study observed no sleep stage effect [12], the other reported that individuals with high dream recall frequency experienced significantly more awakenings from N2 sleep [13]. In contrast, the functional state-shift model [6] argues that dream recall is facilitated when the transition from sleep to wake involves minimal changes in global brain activation, such as awakenings from REM or lighter NREM sleep. In line with this, serial-awakening studies combined with PSG showed that approximately 80% of awakenings from REM sleep result in dream recall, whereas only about 50% of awakenings from NREM sleep do so [18].

Taken together, prior work supports a role for awakenings in dream recall but leaves several fundamental questions unresolved. First, it is unclear whether objective and subjective indices of awakenings reflect the same processes and whether they converge in their predictive power for dream recall. Second, the relative importance of nocturnal awakening frequency versus specific characteristics such as sleep stage, night-half, and duration remains uncertain. Third, prior studies have primarily used retrospective measures to quantify dream recall, single-night laboratory recordings to assess nocturnal awakenings, and group-based comparisons between high- and low-frequency dream recallers to capture between-individual differences in small samples. These methodological approaches limit ecological validity and preclude disentangling stable between-person differences from within-person fluctuations. Moreover, unlike prospective dream diaries, retrospective questionnaires typically do not distinguish between the mere feeling of having dreamed (“white dreams”) and recall of specific content. This distinction is important for understanding the relationship between dream recall and awakenings, as different awakenings may affect these two forms of recall differently: some may increase awareness that a dream occurred without preserving its details, whereas others may help retain specific content. Retrospective measures, therefore, fail to capture these nuances, limiting our understanding of how different awakenings shape different aspects of dream recall.

Here, we address these gaps by leveraging multiple datasets that combine prospective and retrospective measures of dream recall with longitudinal, naturalistic at-home sleep recordings. Using a multilevel framework, we test how awakening frequency, timing (sleep stage and night-half), and duration predict dream recall at both the within-person (state-like) and between-person (trait-like) levels. Finally, in exploratory analyses, we examine the convergence between objective and subjective awakening indices and evaluate whether additional trait- and state-like factors, including memory abilities, substance use, and psychological conditions, are associated with dream recall. Together, this work provides a comprehensive and ecologically grounded account of how nocturnal awakenings and their characteristics shape dream recall, cross-validating across measures, datasets, and levels of analysis.

## Methods

### Study Design and Participants

We conducted our analyses using data from multiple previously collected datasets:

Dataset 1 represents a longitudinal cohort of 905 recruited adults aged 30–39 years. This study was approved by the Institutional Review Board of Radboud University Medical Center (reference number: 2018–4894) in accordance with the latest revision of the Declaration of Helsinki [19], and all participants provided written informed consent. Inclusion criteria were living in the Nijmegen region and willingness and being able to follow the study protocol. Participants were excluded if they had a self-reported history of significant psychiatric or neurological illness, had current brain-affecting diseases, or were using medications such as antidepressants or methylphenidate. Additional exclusion criteria included pregnancy and any contraindications for MRI. Full cohort details are available elsewhere [20]. Assessments took place across three waves spaced ∼4 months apart, over one year. Each wave included multiple nights of home sleep recording and a retrospective dream recall questionnaire.

Dataset 2 consists of 36 adults aged 18–35 years with high dream recall frequency (at least three recalls per week) and at least one lucid dream. This study was approved by METC Oost-Nederland (2014/288, NL45659.091.14) in accordance with the latest revision of the Declaration of Helsinki [19]; all participants provided written informed consent. Participants were excluded if they had a current or past diagnosis of neurological, psychiatric, or neurodegenerative diseases, or had any sleep disorders such as insomnia, sleep apnea, or circadian rhythm disorders. Individuals were also excluded if they had a history of head or brain surgery, epilepsy, pregnancy, or were using psychotropic or sleep medications. Full cohort details are available elsewhere [21]. Assessments comprised nightly home sleep recordings and prospective daily dream diaries for 6 weeks.

Dataset 3 consists of 100 healthy participants aged 18–35 years who had a high dream recall frequency (at least one recall per week), were fluent in English, and were able to sleep in the laboratory setting. This study was approved by CMO Regio Arnhem-Nijmegen (NL75927.091.20); all participants provided written informed consent. Participants were excluded if they had a history of sleep disorders, physical or mental illness, or were taking medications affecting sleep or memory. Additional exclusion criteria included frequent coffee consumption, skin disease at electrode sites, incompatible chronotype, inability to sleep during the adaptation night, MRI contraindications, pregnancy, and irregular sleep patterns prior to experimental sessions. Full cohort details are available elsewhere [22]. Assessments comprised nightly home sleep recordings and prospective daily dream diaries for 4 weeks.

### Data Collection and Analysis

#### Dream Recall

Key outcomes included dream recall frequency (DRF) in Dataset 1, and daily dream recall (white dream [WD] + dream with content [CD] vs. no experience [NE]) and daily content recall (CD vs. WD) in Datasets 2 and 3.

In Dataset 1, DRF was assessed at each wave using the *Mannheim Dream Questionnaire* (MADRE). Participants responded to the item *“How often have you recalled your dreams recently?”* rated on a 0–6 Likert scale, where higher scores indicate more frequent recall [23]. Participants who had not completed at least one DRF questionnaire across the three waves were excluded (n = 27, 3.0%). DRF was treated as a continuous variable.

In Dataset 2, dream recall outcomes were assessed each morning following a recorded night using the item “Do you remember if you dreamed last night? No, I can’t remember having dreamed (NE); yes, I dreamed but I cannot recall a dream (WD); yes, I dreamed and I can recall the dream (CD).” Two binary outcomes were derived: (i) any dream recall, representing the presence of any subjective dream experience (WD+CD vs. NE); and (ii) content recall, representing the level of recall (CD vs. WD). Participants with no completed dream diary were planned to be excluded, however no participants were affected (n = 0, 0.0%). Dream diary entries with missing key fields or filled after another intervening sleep episode were excluded (37 days, 3.0%). Participants who reported having high DRF in the intake session but did not meet the DRF threshold (i.e., at least once a week) based on questionnaires completed afterward were excluded (n=8, 22.2%).

In Dataset 3, dream recall outcomes were assessed each morning following a recorded night using a structured morning diary comprising the items “Did you have any dream, specific thoughts, imagery, sensations, or emotions in the minute before waking up?” (Yes = CD_1 / No), followed, if answered “No,” by “Do you feel as if you had a more detailed dream or specific thoughts, imagery, sensations, or emotions that you have now forgotten?” (Yes = WD_1 / No = NE_1), and “Do you remember any other dreams?” (Yes = CD_2 / No = NE_2). From these, we derived the same two binary outcomes as in Dataset 2: any dream recall (CD_1 or WD_1 or CD_2) and content recall (CD_1 or CD_2 vs. WD_1). Dream diary entries with missing key fields or filled after another intervening sleep episode were excluded (129 days, 5.5%). Participants who reported having high DRF in the intake session but did not meet the DRF threshold (i.e., at least once a week) based on the questionnaires filled out afterward were excluded (n=3, 3.0%). Dream recall and content recall were analyzed as binary variables. The operationalization aligns with established distinctions between the dreaming experience and no experience, and between recall with and without content, as described in prior research (e.g., [24,25]).

#### Main Predictors

In Dataset 1, sleep was recorded in participants’ homes using a portable two-channel EEG headband (ZMax) with frontal derivations (F7–Fpz, F8–Fpz) and integrated tri-axial accelerometry and photoplethysmography sensors. Participants were instructed to apply a disposable hydrogel electrode patch to the forehead and were provided with up to nine recording opportunities (sessions) over seven consecutive nights in each study wave. A given night could include multiple sessions if the device was restarted, whereas some nights could contain no valid recording. We could not independently verify the exact timing and duration of sleep episodes because no sleep diaries were collected, and the episodes lacked corresponding dates. Hence, a set of thresholds was then applied to remove invalid nights or participants. First, participants who had no sleep recordings across the three waves were excluded (n = 61, 7%). Sleep sessions were excluded when total sleep time (TST) was <3 h, to maximize the likelihood of capturing a whole-night recording (2611 recordings, 21%). Although it cannot be confirmed that these sessions always represent individual nights, only eight participants completed eight recorded sleep sessions lasting≥3 h, while the remainder contributed seven or fewer sessions. Sleep staging was performed automatically with Dreamento (stable v1.0; [26]) in 30-s epochs. Signal usability was assessed independently using eegFloss (v1.0; [27,28]) and evaluated in 10-s epochs. Outputs were merged such that a 30-s epoch retained its Dreamento stage if at least 20 s of either frontal channel was usable; otherwise, the epoch was marked unscorable. Stages were labeled as Wake (0), N1 (1), N2 (2), N3 (3), REM (4), or Unscorable (−1). Since not all recorded durations were scorable due to artifacts, sleep sessions were excluded when <75% of epochs were scorable (854 recordings, 9%); all analyses were repeated in sensitivity analyses using a <90% scorable-epoch threshold (2507 recordings, 26%). Participants were excluded if they contributed <3 valid sleep sessions per wave to ensure that the average sleep measures reflect habitual sleep (n=78, 9.8%). Missing data from other waves were left as missing (see Tables S8-S9 for data exclusion flow).

In Datasets 2 and 3, sleep was recorded using Fitbit Inspire 2 devices that infer sleep stages via a fusion of accelerometry and optical photoplethysmography. Participants wore the device nightly throughout the protocol. Main sleep periods were identified using the vendor-defined flag, extracted from JSON exports. When two sleep periods occurred on the same calendar date, and the start of one fell <2 h after the end of the other, they were concatenated into a single session, with the intervening gap coded as Wake. Stages were mapped to a scheme as Wake (0), Light sleep (N1+N2; 2), Deep sleep (N3; 3), and REM (4) through the proprietary Fitbit algorithm. Using the diary-reported wake-up times, we excluded nights when the wearable recording ended more than 30 minutes before the diary-reported wake-up time or when the diary wake-up time was missing (Dataset 2: 112 nights, 9%; Dataset 3: 96 nights, 4.3%). No additional artifact rejection was applied beyond this, and the device’s internal algorithms were used at this stage. Participants were excluded if they contributed <3 valid nights with corresponding morning dream diaries over the study period (Dataset 2: n=0, 0%; Dataset 3: n=1, 1%) (see Table S10-S11 for data exclusion flow).

In all datasets, sleep onset was defined a priori as the first non-Wake epoch within the nocturnal recording [29]. Awakenings were operationalized as contiguous runs of Wake lasting ≥1 epoch (minimum duration 30 s) occurring after sleep onset and within the main sleep period. From the hypnogram, we derived per-night features: total objective awakening count (AW); awakening counts classified by duration (short <1 min, S-AW; medium ≥1 and <2 min, M-AW; long ≥2 min, L-AW); and by preceding sleep stage (REM vs NREM), TST, and final awakening stage (F_REM in Dataset 2 and 3; F_REM% in Dataset 1) and total sleep time (TST, defined as the elapsed time from sleep onset through sleep offset). F_REM% was the proportion of final awakenings from REM, and F_REM was the final awakening stage (contrast coded: REM = 0.5, NREM = −0.5). In Dataset 1, features were averaged within each wave across all valid sessions. In Datasets 2 and 3, features were entered into the models as nightly observations without within-participant averaging. All predictors were treated as continuous variables.

#### Exploratory Predictors

Exploratory predictors were awakening variables occurring in the second half of the night, subjective sleep duration, and subjective awakening count. In Dataset 1, additional exploratory variables were included, which are substance-use, psychological, and cognitive variables. Substance-use variables were alcohol, coffee, and cannabis intake. Psychological variables were state anxiety and depressive symptoms. Memory variables were long-term and short-term associative memory, visual working memory, and auditory working memory.

### Awakening variables in the second half of the night

In all datasets, the main sleep interval was defined as the time between sleep onset and final awakening, as determined from the wearable-derived hypnogram. The midpoint of the sleep period was computed, and duration-stratified awakening counts (short <1 min, S-AW; medium ≥1 and <2 min, M-AW; long ≥2 min, L-AW) were computed for the second half of the night.

### Subjective sleep metrics

In Dataset 1, subjective sleep duration was derived from the *Pittsburgh Sleep Quality Index* (PSQI), which was administered at each assessment wave. Specifically, we used the item: *“During the past month, how many hours of actual sleep did you get at night?”* Responses were coded on a 4-point scale ranging from 0 to 3 (0 = more than 7 hours of sleep; 1 = more than 6 hours of sleep; up to and including 7 hours of sleep; 2 = more than 5 hours of sleep; up to and including 6 hours of sleep; 3 = 5 hours of sleep or less). To align the direction of this measure with the objective sleep metrics (higher values indicating longer sleep), we reverse-coded the item so that higher scores reflect longer subjective sleep duration (0 = 5 hours or less; 1 = more than 5 up to 6 hours; 2 = more than 6 up to 7 hours; 3 = more than 7 hours). The reverse-coded score was analyzed as a continuous measure. Subjective awakening count was derived from the item: *“During the past month, how often have you had trouble sleeping because you wake up in the middle of the night or early morning?”* Responses were coded on a 4-point scale ranging from 0 to 3 (0 = *not during the past month*; 1 = *less than once a week*; 2 = *once or twice a week*; 3 = *three or more times a week*). This variable was analyzed as a continuous measure. In Datasets 2 and 3, participants completed daily sleep diaries each morning, reporting the number of nocturnal awakenings experienced during the preceding night (subjective awakening count), the sleep time, and wake up time. Subjective sleep duration was calculated using sleep time and wake up time. These variables were treated as continuous in the analyses.

### Substance use

In Dataset 1, alcohol, coffee, and cannabis intake were derived from ecological momentary assessments (EMAs) collected during the sleep recording week. For each participant and wave, the proportion of answered beeps with alcohol use, coffee use, and cannabis use was computed and treated as continuous.

### Psychological factors

In Dataset 1, depressive symptoms and state anxiety were assessed with the Inventory of Depressive Symptomatology - Self-Report (IDS-SR) and the State–Trait Anxiety Inventory - State form (STAI-S), respectively. For each instrument, the sum score was computed, following standard scoring guidelines, and treated as continuous.

### Memory

In Dataset 1, associative memory was assessed with a two-alternative forced-choice face–label paired-associates task (short-term test immediately after encoding; long-term test ∼20–30 min later). For each wave, we derived accuracy (proportion correct) on trials for both the short-term and long-term tests. Working memory was measured once and indexed by the visuospatial Corsi (forward) and auditory digit-span (forward) scores, computed according to standard procedures. All memory measures were treated as continuous.

### Control variables

In all datasets, age and sex were treated as continuous predictors. Sex was coded using zero-sum (effects) coding (male = −0.5, female = 0.5). Datasets 2 and 3 were merged into a single dataset for analyses, as both employed identical sleep-recording methods and included individuals with high dream recall frequency. Comparable dream recall outcomes could be derived from both sources. A new continuous variable, dataset, was created to account for dataset-specific effects in subsequent models (Dataset 2 = −0.5, Dataset 3 = 0.5). Another continuous variable, day, was defined as the number of days passed since the diary started.

### Statistical Analysis

#### Variance decomposition of outcomes and predictors

To quantify the proportion of variance attributable to stable between-person differences versus within-person variability in all predictors, we fit unconditional random-intercept mixed-effects models (Outcome/Predictor ∼ 1 + (1|Participant)). Variance components were extracted for the participant-level intercept σ^2^_u0_ and the residual σ^2^_ε_, and the intraclass correlation (ICC) was computed as *ICC = σ^2^_u0_(σ^2^_u0_ + σ^2^_ε_)*

#### Mixed-effects models for dream recall outcomes

We tested the associations between the predictors and DRF (Dataset 1), daily dream recall, and daily content recall (Dataset 2) using separate mixed-effects models. DRF was modelled with linear mixed-effects models (DRF ∼ predictors + age + sex + (1|Participant)); dream recall and content recall with logistic generalized linear mixed-effects models (dream/content recall ∼ predictors + age + sex + dataset + day + (1|Participant)). To disentangle stable individual differences from night-to-night (in Dataset 2+3) or wave-to-wave (in Dataset 1) variations, all time-varying predictors were decomposed into between-person (each participant’s mean across waves or nights) and within-person effects (each participant’s deviation from their mean in each wave or night) and standardized. Hence, coefficients reflect effects per 1 SD change on a standard scale.

Model 1 used total awakening count as a predictor. Model 2 used short, medium, and long awakening counts as predictors. Model 3 used short, medium, and long awakening counts in REM and NREM sleep stages as predictors. Model 4 added TST and F_REM to Model 3 (or F_REM% in Dataset 1). An exploratory model used short, medium, and long awakening counts in REM and NREM sleep stages in the second half of the night, as well as TST and F_REM as predictors (Model E1). The second exploratory model used subjective sleep measures (awakening count and sleep duration) as predictors (Model E2). For DRF, an additional exploratory model used subjective sleep measures, substance-use, psychological, and cognitive variables as predictors (Model E3).

All analyses were performed in R. All hypothesis tests were two-sided with α = 0.05. We reported two-sided P values and 95% two-sided confidence intervals. A sensitivity analysis repeated all models in Dataset 1 under a stricter threshold (≥90% scorable epochs). We considered an effect significant only if it reached significance in both the primary and sensitivity analyses, and we reported both P values (main, *P*; sensitivity analysis, *P*_*s*_) only when they differed in whether they met the significance threshold.

Although the DRF item is ordinal, we treated the 0–6 score as approximately continuous and analyzed it with linear mixed-effects models because our aim was to estimate overall graded associations with dream recall frequency. Ordinal mixed-effects models would also have been possible, but they require additional assumptions and would add complexity to an already detailed repeated-measures analysis, especially given the relatively sparse observations in some response categories at the ends of the scale. Linear mixed-effects models therefore provided a simpler, more interpretable, and literature-comparable framework.

We did not apply a blanket correction for multiple comparisons. Our models were specified a priori and interpreted individually across datasets, and the analyses were conducted in a mixed-model framework that jointly estimates related effects. This choice follows arguments against routine multiplicity adjustment for all analyses and aligns with work showing that hierarchical models can partially address multiplicity through partial pooling [30,31]. We note, however, that preregistration alone does not automatically remove multiplicity concerns; accordingly, we report all tests transparently and interpret marginal findings cautiously.

### Deviations from preregistration

This study was preregistered at OSF prior to analyses: [link]. We made four changes from the preregistered plan. First, rather than analyzing the two Fitbit cohorts separately, we pooled data from Datasets 2 and 3 and included a fixed effect for dataset. This choice was made because the same device and closely aligned diary instruments were used across studies, intended to increase sample size and generalizability while accounting for cohort differences. Second, owing to collinearity between the awakening counts and wake after sleep onset (WASO; up to r = 0.7), WASO was excluded from the primary models. Third, for time-varying predictors, we applied a within–between decomposition, rather than analyses based on raw variables, to better isolate within-person effects. Lastly, we additionally controlled for TST in the last preregistered model to account for the effect of sleep duration.

## Results

### Sample characteristics and data structure

Dataset 1 was used for analyses of DRF (see Fig. 2a for study design). 708 participants were included (58.5% female; mean age 33.8 ± 2.8 years; *all-recaller* cohort), yielding 1,477 wave-level observations (wave 1, n = 600; wave 2, n = 446; wave 3, n = 431). Participants contributed 3–8 usable nights per wave (mean nights per participant per wave 5.0 ± 1.4).

**Fig. 2:**
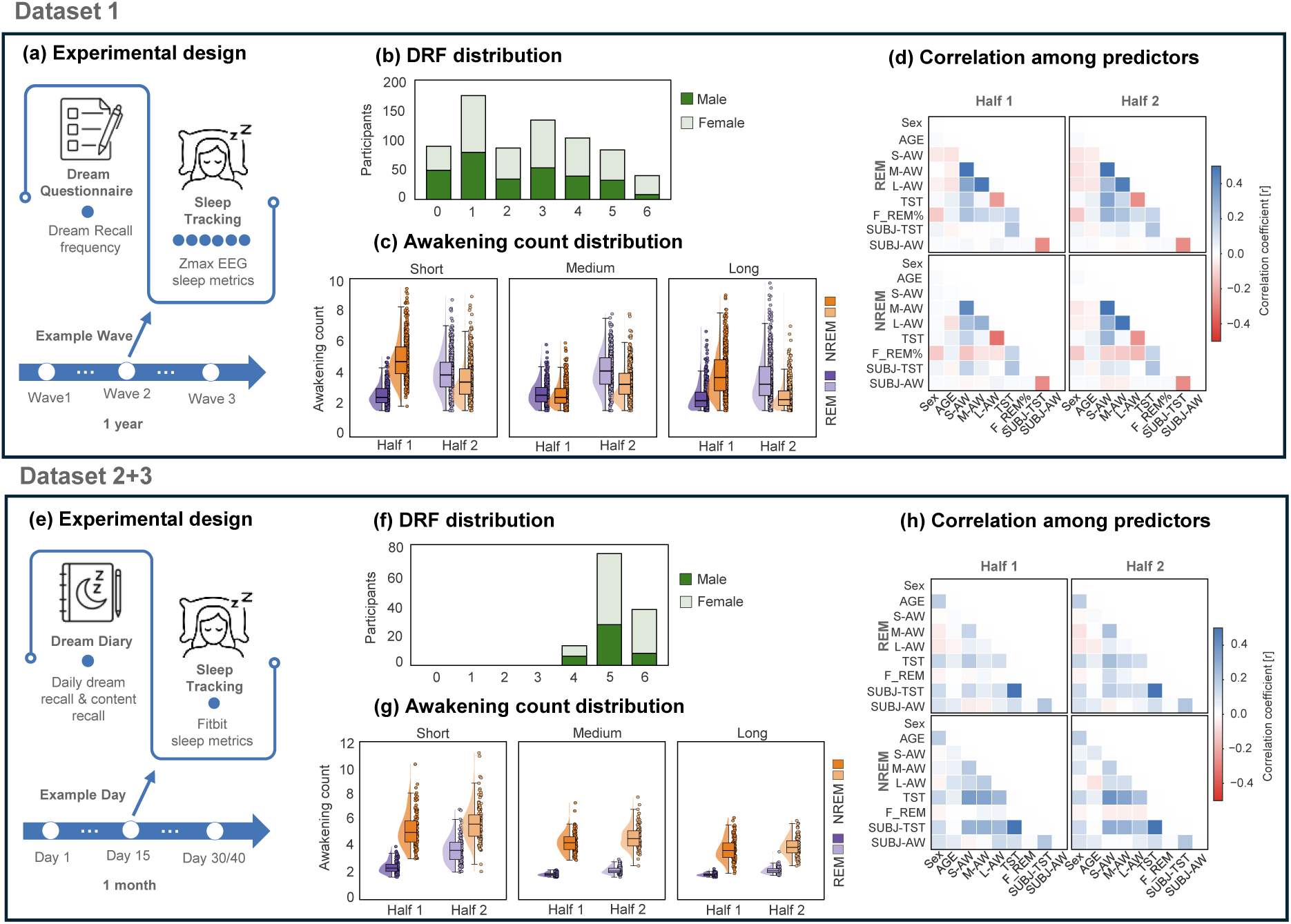
Description of the experimental design and the collected data **a-d**, Dataset 1; **e-h**, Dataset 2+3. **(a,e)** Outline of the experimental design. **(b,f)** Dream recall frequency (DRF) distribution by sex; Colors indicate sex: light green, female; dark green, male; Dream recall frequency (0–6): 0 = Never; 1 = Less than once a month; 2 = About once a month; 3 = Two or three times a month; 4 = About once a week; 5 = Several times a week; 6 = Almost every morning. **(c,g)** Awakening count distributions for short (<1 min), medium (≥1 and <2 min), and long (≥2 min) awakenings from REM (dark purple/light purple) and NREM (dark orange/light orange) in the first and second half of the night (Half 1, Half 2); Dots represent participant means. **(d,h)** Correlations (pairwise Pearson’s r) among objective and subjective awakening and sleep measures; Color intensity indicates correlation magnitude and sign (red, negative; white, ∼0; blue, positive). Abbreviations: TST, total sleep time; S-AW/M-AW/L-AW, short/medium/long awakening count; F_REM (%), (percentage of) final awakenings preceded by REM; REM/NREM, sleep stage; Half1/Half2, first/second half of the night; SUBJ-TST, subjective sleep duration; SUBJ-AW, subjective awakening count.

Datasets 2 and 3 were pooled for analyses of daily dream recall and content recall (see Fig. 2e for study design). In total, 124 participants were included (66.9% female; mean age 24.4 ± 3.6 years; *high-recaller* cohort), yielding 2,896 night-level observations. Participants contributed 3–42 usable nights (mean nights per participant 23.4 ± 9.2).

### Dream Recall: Descriptive Statistics

In Dataset 1, participants’ self-reported DRF on the MADRE 0–6 scale averaged 2.74 ± 1.71 when pooled across all participant–wave observations, meaning on average across all time points, participants remembered dreams “about once a month” to “two or three times a month” (see Fig. 2b and Table 1 for the full distribution). Female participants reported slightly higher DRF than male participants (males, mean = 2.52; females, mean = 2.86; Wilcoxon rank-sum test, *P < 0.001*). ICC indicated that 74% of the variance in DRF was attributable to between-person differences (26% within-person variability over waves).

In Dataset 2+3, participants’ self-reported DRF on the MADRE 0–6 scale averaged 5.19 ± 0.61, meaning they remembered their dreams “several times per week” to “almost every morning” (see Fig. 2f and Table 2 for the full distribution). Females showed slightly higher questionnaire DRF than males (males, mean = 4.97; females, mean = 5.26; Wilcoxon rank-sum test*, P < 0.001*). Prospective-based DRF was derived from prospective diary measures and converted to the MADRE scale to allow comparison with retrospective questionnaire-based DRF. Prospective-based DRF calculated from the dream recall measure (WD + CD; 4.90 ± 1.10) differed significantly from questionnaire-based DRF (Wilcoxon signed-rank test, *P* = 0.03) and showed a moderate positive association with it (Spearman’s correlation, *r* = 0.38, *P* < 0.001). Prospective-based DRF derived from the content recall measure (CD; 3.54 ± 1.30) showed a similar pattern (*P* < 0.001; *r* = 0.40, *P* < 0.001). Across all morning reports, 18.3 % of morning reports indicated NE, 58.1% indicated CD, and 23.5% indicated WD. Female participants showed a higher daily probability of WD (mean difference 0.09, *P = 0.01*) and CD (mean difference = 0.11, *P < 0.01*). ICC indicated substantial within-person variability in both dream recall (CD+WD vs. NE) and content recall (CD vs. WD). For dream recall, 34% of the variance was attributable to stable between-participant differences (66% within-person variability over days). For content recall, 26% of the variance was attributable to stable between-participant differences (74% within-person variability over days).

### Predictors: Descriptive Statistics

In dataset 1, ICC indicated a considerable within-person variability over waves in the main predictors (40–70%; see Table 3 for full descriptive details). Exploratory predictors showed mixed within-person variability over waves (21-78%; see Table 4 for complete descriptive details). Awakening counts showed clear stage-, duration-, and night-half-dependent distributions, with NREM awakenings being more frequent in the first half of the night and REM awakenings in the second half of the night (see Fig. 2c). Correlations among main predictors are shown in Fig. 2d, and detailed between-person and within-person correlation matrices are shown in Fig. S1.

In Dataset 2+3, ICC indicated a substantial within-person variability over nights in the main and in the exploratory predictors (63-93%; see Table 5 and Table 6 for full descriptive details). Awakenings showed clear stage-, duration-, and night-half-dependent distributions, with NREM awakenings being more frequent than REM awakenings overall (see Fig. 2g). Correlations among main predictors are shown in Fig. 2h and detailed between-person and within-person correlation matrices are shown in Fig. S2.

### Mixed-Effects Models for dream recall outcomes

#### Dream Recall Frequency in Dataset 1 (all-recaller cohort)

In this section, we report between-person (habitual) and within-person (wave-to-wave) associations between primary and exploratory predictors and DRF in the main (Models 1–4; Tables 7-10, sensitivity analyses; Tables S1-S7) and exploratory models (Models E1–E3; Tables 11-13).

### Between-person associations (habitual)

Habitual awakening count was positively associated with DRF in the all-recaller cohort (β = 0.14, 95% CI 0.02–0.25; *P = 0.02*; Model 1). When it was decomposed by duration, only habitual long awakening count was positively associated with DRF (β = 0.20, 95% CI 0.04–0.35; *P = 0.01*), with no significant effects for habitual short or medium awakening counts (*P ≥ 0.18*; Model 2). Stratifying awakening counts of different lengths by preceding sleep stage showed that habitual short awakening count in NREM was positively associated with DRF (β = 0.17, 95% CI 0.01–0.32; *P = 0.04*), whereas the effect of habitual long awakening count did not meet the significance criterion (*P < 0.01*; *P*_*s*_ *= 0.06*; Model 3). Notably, the effect of habitual short awakening count in NREM was not robust to adjustment for TST and F_REM% (β = −0.0; *P = 0.9*); after this adjustment, habitual long awakening count in REM was again positively associated with DRF (β = 0.27, 95% CI 0.09–0.44; *P < 0.01*), with no independent contributions of TST and F_REM% (*P ≥ 0.05*; Model 4). This pattern continued to persist in the second-night-half exploratory model (Model E1; Fig. 3a).

**Fig. 3:**
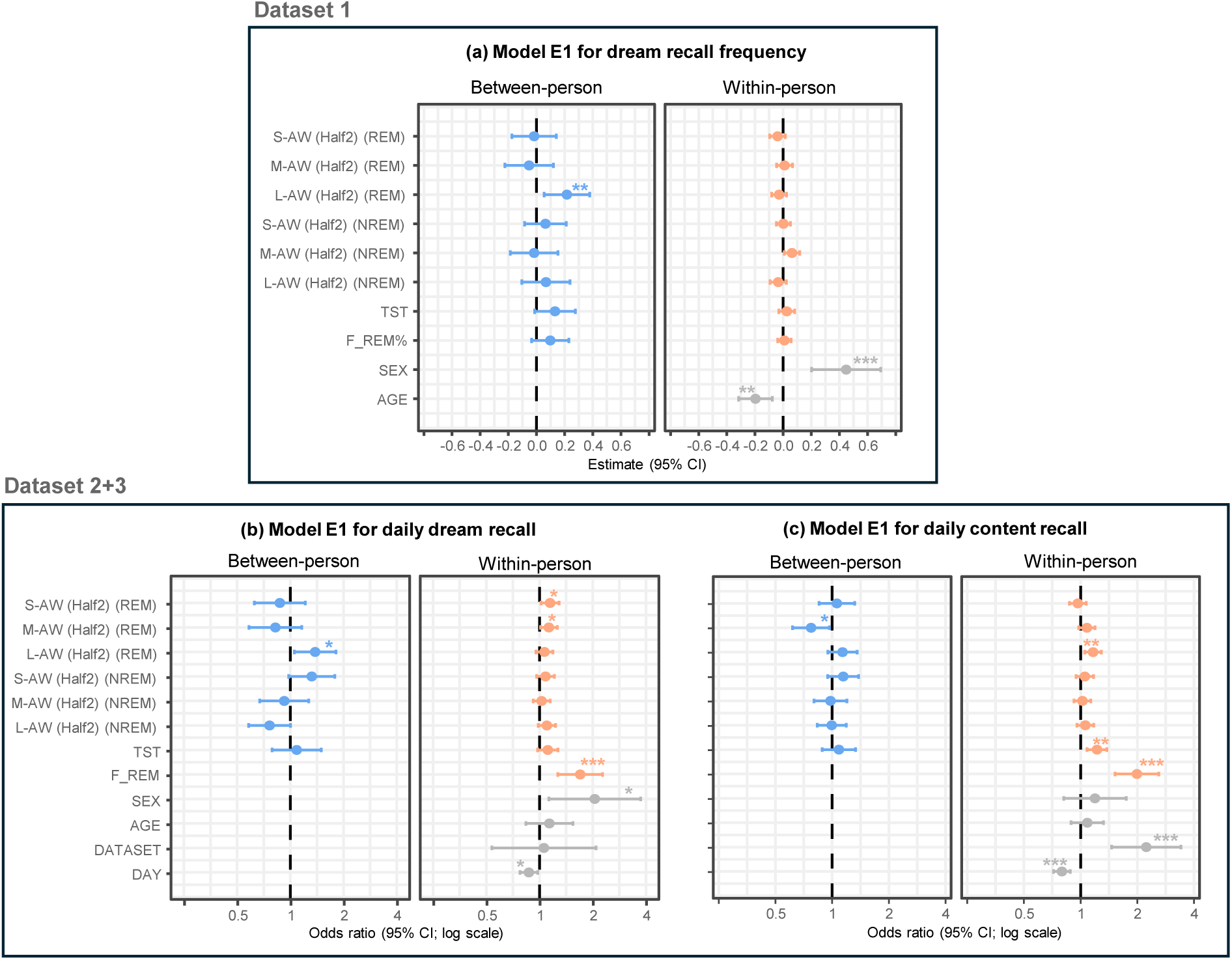
Predictors of dream recall outcomes Mixed-effects models predicting **(a,b)** dream recall frequency (DRF), **(c,d)** dream recall (WD+CD vs. NE), **(e,f)** dream content recall (CD vs. WD) from objective awakening measures across restricted to the second half of the night (a,b.c; Model E1). Predictors were decomposed into between-person components (person-level means; blue) and within-person components (person-mean–centered deviations; orange); non-decomposed covariates are shown in grey. Estimates represent the expected change in outcome per 1 SD increase in each predictor. Odds ratios show the expected change in the odds of outcome per 1 SD increase in each predictor. Sex was contrast-coded (−0.5 = male, +0.5 = female). Asterisks indicate statistical significance: *p < .05, **p < .01, ***p < .001. Abbreviations: DAY, study day; TST, total sleep time; S-AW/M-AW/L-AW, short/medium/long awakening count; F_REM (%), final awakenings from REM (%); REM/NREM, preceding sleep stage.

The exploratory model including subjective sleep measures showed that both habitual awakening count (β = 0.33, 95% CI 0.21–0.46; *P < 0.001*) and habitual TST (β = 0.15, 95% CI 0.03–0.28; *P = 0.02*) were positively associated with DRF (Model E2). When we added substance-use, cognitive and psychological exploratory variables to this model, only habitual awakening count (β = 0.33, 95% CI 0.16–0.50; *P < 0.001*) and habitual cannabis intake (β = 0.19, 95% CI 0.05–0.33; *P < 0.01*) were positively associated with DRF, whereas the remaining variables were not significantly associated with DRF (*P ≥ 0.16*; Model E3).

### Within-person associations (wave-to-wave)

Within-person deviations from habitual awakening count were not associated with DRF (β = 0.00, 95% CI −0.04 to 0.05; *P = 0.91*; Model 1). When they were decomposed by duration, lower-than-usual long awakening count was associated with higher DRF (β = −0.06, 95% CI −0.12 to −0.01; *P* = 0.03), whereas within-person deviations in short and medium awakening counts did not meet the significance criterion (short, *P* = 0.81; medium, *P = 0.04*; *P*_*s*_ *= 0.06;* Model 2). The small negative effect of within-person deviations in long awakening count did not persist after further stratifying awakening counts by sleep stage (*P* = 0.14), while other effects remained non-significant (*P* ≥ 0.1; Model 3). This null pattern continued to persist after additional adjustment for TST and F_REM% (Model 4), and in the second-night-half exploratory model (Model E1, Fig. 3a), with no independent contributions of TST and F_REM% (*P* ≥ 0.1).

The exploratory model, including subjective sleep measures, showed that within-person deviations in awakening count and TST were not associated with DRF (*P* ≥ 0.4; Model E2). Finally, the last exploratory model showed that within-person deviations in none of the exploratory predictors were significantly associated with DRF (*P* ≥ 0.63; Model E2).

### Covariates and explained variance

Across the main and exploratory models, covariates showed consistent patterns: female sex was associated with higher DRF (β = 0.39–0.47; *P* ≤ 0.01, not significant in EM3, *P = 0.21*), whereas age was associated with lower DRF (β = −0.17 to −0.21; *P* ≤ 0.01). Explained variance increased only modestly with model refinement (Table 14). Although the final main model captured substantial variance overall, variance attributable to fixed effects remained small (Model 4; marginal R² = 0.05, conditional R² = 0.72). The final exploratory model captured more overall variance (EM3; marginal R² = 0.08, conditional R² = 0.74), albeit in a slightly smaller subsample.

### Daily dream recall in Dataset 2+3 (high-recaller cohort)

In this section, we report between-person (habitual) and within-person (night-to-night) associations between primary and exploratory predictors and daily dream recall in the main models (Models 1-4; Tables 15-18) and in the exploratory models (Models E1-E2; Tables 19-20).

### Between-person associations (habitual)

Habitual awakening count was not associated with dream recall (WD+CD vs. NE) in the high-recaller cohort (*P* = 0.57; Model 1), and decomposing it by duration likewise yielded no significant associations for habitual short, medium, or long awakening counts (all *P* ≥ 0.17; Model 2). When habitual duration-stratified awakening counts were further stratified by sleep stage, higher habitual long awakening count in REM (OR = 1.38, 95% CI 1.05–1.80; *P* = 0.02), higher habitual short awakening count in NREM (OR = 1.46, 95% CI 1.08–1.99; *P* = 0.02), and lower habitual long awakening count in NREM (OR = 0.68, 95% CI 0.51–0.91; *P* < 0.01) were associated with higher odds of dream recall (Model 3). These associations were unchanged after controlling for TST and F_REM, with no independent contribution of TST (*P = 0.3*; Model 4). In the second-night-half exploratory model, only higher long awakening count in REM was associated with higher odds of dream recall (OR = 1.38, 95% CI 1.05–1.81; *P* = 0.02; Model E1; Fig. 3b).

The exploratory model incorporating subjective sleep measures showed that higher habitual awakening count predicted higher odds of dream recall (OR = 1.38, 95% CI 1.04–1.84; *P* = 0.03), with no independent contribution of habitual TST (*P=0.9*; Model E2).

### Within-person associations (night-to-night)

Higher-than-usual awakening count was associated with higher odds of dream recall (OR = 1.28, 95% CI 1.15–1.42; *P* < 0.001; Model 1). When the awakening count was decomposed by duration, only the higher-than-usual short awakening count was associated with higher odds of dream recall (OR = 1.22, 95% CI 1.09–1.36; *P* < 0.001), with no independent contribution of within-person deviations in medium or long awakening counts (*P* ≥ 0.13; Model 2). When awakening counts were stratified by sleep stage, only a higher-than-usual short awakening count in REM was associated with higher odds of dream recall (OR = 1.23, 95% CI 1.10–1.38; *P* < 0.001, Model 3). This pattern persisted after adjustment for TST and F_REM and in the second-half-of-night exploratory model, with additional positive associations for F_REM (OR = 1.69, 95% CI 1.26–2.27; *P* < 0.001; Model 4) and medium awakening count in REM (OR = 1.13, 95% CI 1.01–1.26; *P* = 0.03; Model E1, Fig. 3b).

In the last exploratory model incorporating subjective sleep measures, higher-than-usual awakening count (OR = 1.30, 95% CI 1.15–1.47; *P* < 0.001) and longer-than-usual TST (OR = 1.30, 95% CI 1.16–1.44; *P* < 0.001) were significantly associated with dream recall (Model E2).

### Covariates and explained variance

Across the main models and exploratory models, the covariates contributed consistent effects: female sex was associated with higher odds of daily dream recall (OR = 1.92–2.29; *P* ≤ 0.02; not significant in E2, *P = 0.05*), study day was negatively associated with dream recall (OR = 0.87; *P* ≤ 0.02), whereas age and dataset showed no reliable associations (all *P* ≥ 0.20). Explained variance increased only modestly with successive model refinements (Table 21). The final main model captured limited variance overall, and inclusion of random effects added little beyond the fixed predictors (Model 4; marginal R² = 0.07, conditional R² = 0.11). Restricting objective predictors to the second half of the night (EM1; marginal R² = 0.06, conditional R² = 0.10), and replacing objective sleep measures with subjective ones (EM2; marginal R² = 0.05, conditional R² = 0.08) slightly reduced the explained variance.

### Daily content recall in Dataset 2+3 (high-recaller cohort)

In this section, we report between-person (habitual) and within-person (night-to-night) associations between primary and exploratory predictors and daily content recall in the main models (Models 1-4; Tables 22-25) and in the exploratory models (Models E1-E2; Tables 26-27).

### Between-person associations (habitual)

Habitual awakening count was not associated with content recall (CD vs. WD) in the high-recaller cohort (OR = 1.06, 95% CI 0.90–1.25; *P = 0.49*; Model 1), and decomposing it by duration similarly yielded no significant associations for short, medium, or long awakening count (OR = 0.94–1.12; all *P ≥ 0.24*; Model 2). When habitual duration-stratified awakening counts were further stratified by sleep stage, only a lower habitual medium awakening count in REM was associated with higher odds of content recall (OR = 0.72, 95% CI 0.56–0.92; *P < 0.01*; Model 3). This pattern continued to persist after controlling for TST and F_REM (Model 4) and in the second-night-half exploratory model (Model E1; Fig. 3c).

In the exploratory model incorporating subjective sleep measures, habitual awakening count and habitual TST were not significantly associated with content recall (P ≥ 0.10; Model E2).

### Within-person associations (night-to-night)

Higher-than-usual awakening count predicted higher odds of content recall (OR = 1.25, 95% CI 1.13–1.38; *P* < 0.001; Model 1). After decomposing awakening count by duration, higher-than-usual short awakening count (OR = 1.14, 95% CI 1.04–1.26; *P* < 0.01) and long awakening count (OR = 1.12, 95% CI 1.02–1.24; *P* = 0.02) predicted higher odds of content recall, with no independent effect of medium awakening count (OR = 1.10, 95% CI 0.99–1.21; *P* = 0.07; Model 2). In the stage-stratified model, higher-than-usual long awakening count in REM (OR = 1.14, 95% CI 1.04–1.26; *P* < 0.01) and higher-than-usual short awakening count in NREM (OR = 1.16, 95% CI 1.05–1.28; *P* < 0.01) predicted higher odds of content recall (Model 3). However, after adjusting for TST and F_REM, only the effect of within-person deviations in long awakening count in REM (OR = 1.12, 95% CI 1.01–1.24; *P* = 0.03) remained significant, with additional positive associations for longer-than-usual TST (OR = 1.22, 95% CI 1.06–1.40; *P* < 0.01), and F_REM (OR = 1.98, 95% CI 1.52–2.58; *P* < 0.001; Model 4). This pattern continued to persist in the second-half-of-night exploratory model (Model E1; Fig. 3c).

In the last exploratory model incorporating subjective sleep measures, longer-than-usual TST (OR = 1.31, 95% CI 1.18–1.45; *P* < 0.001) was associated with higher odds of content recall, with no independent effect of within-person deviations in awakening count (P = 0.43; Model E2).

### Covariates and explained variance

Across the main models and exploratory models, covariates showed stable effects: dataset (being in Dataset 3) was strongly associated with higher odds of content recall (OR = 2.22–2.26; *P < 0.001*), and study day was consistently associated with lower odds of content recall (OR = 0.79–0.80; *P < 0.001*). By contrast, sex and age were not reliably associated with content recall (all P ≥ 0.16). Explained variance increased only modestly with successive model refinements (Table 28). The final main model captured limited variance overall, and inclusion of random effects added little beyond the fixed predictors (Model 4; marginal R² = 0.08, conditional R² = 0.12). Restricting objective predictors to the second half of the night slightly increased the explained variance (EM1; marginal R² = 0.09, conditional R² = 0.12). In contrast, replacing objective sleep measures with subjective ones reduced the explained variance (EM2; marginal R² = 0.06, conditional R² = 0.09).

## Discussion

Our study provides converging support for the arousal–retrieval model [5] and the functional state-shift model [6], and extends previous findings [12,13] by revealing additional nuances in the trait-like awakening profile associated with high dream recall frequency: a higher propensity for longer REM awakenings, reflected by more habitual long REM awakenings and/or fewer habitual medium REM awakenings, and a higher propensity for shorter NREM awakenings, reflected by more habitual short NREM awakenings and/or fewer habitual long NREM awakenings. Specifically, in the all-recaller cohort, more habitual long REM and short NREM awakenings were associated with higher dream recall frequency. In the high-recaller cohort, more habitual long REM and short NREM awakenings, together with fewer habitual long NREM awakenings, predicted daily dream recall, whereas fewer habitual medium REM awakenings predicted daily content recall. Notably, NREM awakening effects survived adjustment for sleep duration only in the high-recaller cohort, whereas REM awakening effects remained robust after adjustment for sleep duration in both cohorts. This suggests that the higher frequency of long REM awakenings in individuals with high dream recall frequency is not merely explained by longer sleep providing more opportunities to awaken [32–34], but may instead reflect a partially independent trait. This greater tendency to awaken during the night, independent of sleep duration, may also be partly explained by increased stimulus reactivity during sleep, as reported in individuals with high dream recall frequency [11,12].

The night-half analyses further suggest that habitual REM and NREM awakenings may contribute through different pathways. REM effects remained evident when awakenings were restricted to the second half of the night, consistent with the idea that long REM awakenings near the final awakening and morning report may directly support dream encoding and increase the likelihood that dream memories remain accessible for recall. By contrast, NREM effects were not evident when awakenings were restricted to the second half of the night, suggesting that they may be driven primarily by earlier-night awakenings or broader sleep-architecture features rather than by awakenings near morning recall. More specifically, having more habitual short NREM awakenings and fewer habitual long NREM awakenings may benefit daily dream recall in a less direct way. Long NREM awakenings earlier in the night may promote deeper sleep later in the night by increasing homeostatic sleep pressure, and awakenings from deeper NREM sleep later in the night may be less optimal for dream encoding and retrieval due to sleep inertia and more gradual awakening processes [35,36]. This interpretation is consistent with a recent study reporting a negative association between longer deep sleep and daily dream recall [25], and with the positive correlation we observed between long NREM awakenings earlier in the night and deep sleep duration later in the night (Fig. S2a). However, because we did not formally test mediation, this interpretation remains tentative.

Beyond these trait-like differences, our results also highlight important within-person dynamics. Within-person changes in awakening measures across the three time points (over a year) were not associated with questionnaire-based dream recall frequency in the all-recaller cohort. This may reflect the limited sensitivity of wave-level questionnaire measures and sleep metrics averaged over a week for tracking within-person variation, given that dream recall frequency has been shown to be relatively stable across years [37]. Although dream recall frequency assessed over broader time windows may remain relatively stable, meaningful day-to-day variation may still occur, as observed in our study. In the high-recaller cohort, within-person changes in awakening measures showed a clear relationship with daily dream recall outcomes, with the most robust effects emerging for REM awakenings: more long REM awakenings (≥2 minutes) than usual increased the likelihood of remembering dream content, whereas more short and medium REM awakenings (<2 minutes) than usual increased the likelihood of reporting a sense of having dreamed. These within-person effects were independent of within-person changes in sleep duration and final awakening from REM sleep, and were primarily driven by awakenings occurring in the second half of the night. Thus, our study shows for the first time that having more long awakenings is not only a trait, as shown in previous studies [12,13], but also influences dream recall on the state level, and that it predicts recall of specific dream content in individuals with high dream recall frequency.

These habitual and night-to-night patterns were also partly supported by subjective measures. Reported habitual awakenings predicted both dream recall frequency and daily dream recall, and more subjectively reported awakenings than usual increased the likelihood of reporting a sense of having dreamed. However, subjective awakenings appeared less specific for distinguishing content recall from the mere sense of having dreamed. This may reflect the lower specificity of subjective awakening measures: subjective awakenings capture whether participants noticed and remembered waking during the night, but they do not distinguish awakenings by sleep stage, duration, or timing. They may therefore reflect a general tendency toward perceived sleep fragmentation, which is sufficient to increase the likelihood of reporting that a dream occurred, but not specific enough to predict whether dream content was retained.

Final awakening from REM sleep also predicted both daily dream recall and daily content recall, with a stronger effect than any of the other observed predictors, further confirming that timing and sleep stage of an awakening are important determinants of dream recall. Although we included both last-minute dreams and earlier dreams in our analysis, the effect of final awakening from REM sleep might be due to last-minute dreams being much more commonly reported. Independent of awakening-related effects, within-person increases in objective sleep duration predicted only content recall in the high-recaller cohort. This association may reflect either the beneficial effects of longer sleep on memory processes or simply the greater opportunity for more and longer dreams. By contrast, within-person increases in subjective sleep duration were associated with both daily dream recall and daily content recall. This partial divergence between subjective and objective measures may be due to the fact that, in the subjective models, although awakening effects were controlled, the effect of a final awakening from REM sleep was not; therefore, longer reported sleep may partly reflect an increased likelihood of waking from REM in the morning.

In addition to the primary findings, several secondary results emerged for control variables and other exploratory predictors. While these findings should be interpreted cautiously, they provide further context for understanding variation in dream recall.

With regard to demographic factors, some previous studies have reported a small negative effect of age and a positive effect of female sex on dream recall [33,38–40]. In our study, age predicted dream recall frequency in the all-recaller cohort, but not daily dream recall measures in the high-recaller cohort. One possible explanation for this difference is the age distribution of the two cohorts. Participants in the all-recaller cohort were adults in their fourth decade of life, whereas participants in the high-recaller cohort were in their third decade. It may therefore be that the age effect is less apparent at younger ages. We also found that female sex predicted dream recall frequency in the all-recaller cohort and daily dream recall in the high-recaller cohort, but not daily content recall in the high-recaller cohort. This pattern may suggest that female participants were more likely to report the sense of having dreamed, rather than showing better retention or retrieval of dream details once recalled, and is consistent with findings indicating that the association may depend on how dream recall is measured [41].

Beyond demographic variables, some of the secondary findings point to an important role of methodological context in shaping recall outcomes. In line with a previous study showing that keeping a diary negatively affected dream recall in individuals with high dream recall frequency [41], we found that it not only reduced daily dream recall but also reduced the odds of daily content recall, suggesting that motivation may affect both measures similarly. We also observed that being in Dataset 3 (versus Dataset 2) had a positive effect on content recall, possibly due to differences in the type of diaries used in these datasets, including the wording of the questions (e.g., “having any dreams” vs. “any thought, sensation, or emotion”) and the time period they referred to (the 1 minute before awakening and the rest of the night vs. the whole night; see Methods for details).

By contrast, several psychological and cognitive variables discussed in prior work on dream recall [42,43] did not show convincing associations in our sample. We did not provide convincing evidence for effects of either habitual levels or within-person deviations in memory abilities, including visual and auditory working memory, and short- and long-term associative memory, nor for effects of non-clinical anxiety or depression in our cohort. These null findings should, however, be interpreted with caution. In the case of depression and anxiety, scores in our sample remained below the clinical cut-offs (see Table 4), which may have limited the ability to detect meaningful associations. In the case of memory, the relevant domain may simply not have been captured by the measures included, as potentially more informative constructs, such as autobiographical memory, were not assessed. It is also possible that memory-related effects on dream recall are more likely to emerge when recall is characterized in greater detail, both quantitatively (how much is remembered) and qualitatively (e.g., vividness or richness of recall).

Finally, one exploratory finding that remained after adjustment for a broad set of covariates concerned cannabis use. Even after controlling for age, sex, sleep, and awakening variables, anxiety, depression, and alcohol and caffeine use, habitual cannabis intake showed a small positive association with dream recall frequency. This suggests that the relationship is not fully explained by affective symptoms, common co-substance use, or overall sleep fragmentation. Rather, it may reflect phenomenological differences in dreams that facilitate encoding, or intermittent withdrawal associated with subtle REM rebound effects [44].

Although our results identify several significant predictors of dream recall, they also indicate that much of the relevant variance remains unexplained. This may partly reflect limitations of the current study. First, we captured recall primarily through broad measures of whether any dream was remembered. Future studies should use diaries and questionnaires that assess both the quantity and quality of dream reports, which may be better suited to studying the effects of nocturnal awakenings on dream recall. Second, we conducted the daily recall analyses only in a high-recaller cohort, leaving open the question of whether stronger associations would be observed in samples with greater variability in dream recall frequency. Third, we used both Fitbit and ZMax devices and obtained broadly similar results across the two systems; however, both devices have known hardware- and algorithm-related shortcomings that should be considered when interpreting the findings. Fitbit infers sleep stages and awakenings indirectly from movement and cardiac signals and may miss brief awakenings or misclassify stage transitions, whereas ZMax, although EEG-based, remains limited by its frontal montage, signal quality variability, and automated scoring constraints relative to full PSG. Future studies would therefore benefit from improved signal-quality control and validation against concurrent gold-standard polysomnography to better quantify device-specific bias and strengthen confidence in sleep stage and awakening estimates.

The modest explained variance may also be informative at a conceptual level. Specifically, it supports the idea that awakenings may provide opportunities for dream recall, but that additional factors likely determine whether these opportunities lead to successful recall. One such factor may be the phenomenological features of dreams, which future studies should examine alongside awakening and sleep metrics. This possibility is particularly interesting because certain dream characteristics may themselves promote more abrupt awakenings, especially from REM sleep. Future research should therefore aim to disentangle these potentially bidirectional pathways.

Taken together, our findings indicate that dream recall is shaped not only by how often individuals awaken during the night but also by when those awakenings occur, the sleep stage from which they arise, and how long they last. By examining both between-person and within-person sources of variation, our results further suggest that some awakening-related measures associated with dream recall at the within-person level are mirrored at the between-person level, whereas others remain specific to the within-person level among individuals with high dream recall frequency. Moreover, we show that REM and NREM awakenings may contribute to dream recall through partly distinct pathways, and that awakenings of different durations contribute to different levels of dream recall. Overall, this study offers a clearer framework for understanding how nocturnal awakenings and their characteristics shape dream recall by refining the arousal–retrieval model [5] and the functional state-shift model [6] and confirming their ecological validity.

## Data availability

The data analyzed in this study were obtained in part from the Healthy Brain Study cohort. Requests to access Healthy Brain Study data should be directed to the Healthy Brain Study via its data-access procedure (link). Data from the other contributing datasets are on the Donders Data Repository and available upon request from Martin Dresler.

## Acknowledgments

This work was supported by the German Academic Exchange Service (DAAD, Research Grants – Doctoral Programmes in Germany, 2021/22, to S.A.), the Swiss National Science Foundation (P2ZHP1_195248, P500PS_210884, and P5R5PS_230534, to S.F.S.), the Cogito Foundation (20_123-S, to S.F.S. and M.D.), the DreamScience Foundation and the International Association for the Study of Dreams (IASD; to S.F.S. and M.D.), and the Dutch Research Council (NWO; 016.Vidi.185.142, to M.D.).

This research was conducted using the Healthy Brain Study resource under Release Number 2024-4. The authors wish to acknowledge the services of the Healthy Brain Study team and the contribution of the Healthy Brain Study consortium. We also thank the researchers, students, and technical staff who assisted with the collection of the other datasets included in this work, as well as all study participants.

## Author contributions

**M.D.:** Conceptualization, Supervision, Funding acquisition, Resources, and Writing – review & editing. **M.J.E.:** Data curation and Writing – review & editing. **N.A.:** Conceptualization, Supervision, and Writing – review & editing. **S.A.:** Conceptualization, Methodology, Data curation, Formal analysis, Funding acquisition, Investigation, Software, Visualization, Writing – original draft, and Writing – review & editing. **S.F.S.:** Conceptualization, Methodology, Data curation, Funding acquisition, Supervision, and Writing – review & editing. All authors reviewed and approved the final manuscript.

## Competing interests

The authors declare no competing interests.

### Dream Recall: Descriptive Statistics

**Table 1.** DRF distribution in Dataset 1.

| Statistic | Wave 1 | Wave 2 | Wave 3 |
| --- | --- | --- | --- |
| Participants (n) | 600 | 446 | 431 |
| Mean ± SD | 2.83 ± 1.77 | 2.67 ± 1.69 | 2.6 ± 1.75 |
| Never (0) | 64 (10.7%) | 45 (10.1%) | 49 (11.4%) |
| Less than once a month (1) | 112 (18.7%) | 99 (22.2%) | 102 (23.7%) |
| About once a month (2) | 80 (13.3%) | 53 (11.9%) | 62 (14.4%) |
| Two or three times a month (3) | 110 (18.3%) | 101 (22.6%) | 79 (18.3%) |
| About once a week (4) | 112 (18.7%) | 77 (17.3%) | 60 (13.9%) |
| Several times a week (5) | 85 (14.2%) | 50 (11.2%) | 58 (13.5%) |
| Almost every morning (6) | 37 (6.2%) | 21 (4.7%) | 21 (4.9%) |

Counts represent unique participant–wave observations; percentages are within wave.

**Table 2.** DRF distribution in Dataset 2+3.

| Statistic | Value |
| --- | --- |
| Participants (N) | 124 |
| Range | 4–6 |
| Mean ± SD | 5.19 ± 0.61 |
| Never (0) | 0 (0.0%) |
| Less than once a month (1) | 0 (0.0%) |
| About once a month (2) | 0 (0.0%) |
| Two or three times a month (3) | 0 (0.0%) |
| About once a week (4) | 13 (10.5%) |
| Several times a week (5) | 74 (59.7%) |
| Almost every morning (6) | 37 (29.8%) |

Note. Counts are unique participants; percentages are within dataset.

### Predictor Variables: Descriptive Statistics

**Table 3.** Descriptive statistics of main predictors in Dataset 1.

| Variable | N (obs) | N (ID) | Mean ± SD | Range | Between % | Within % |
| --- | --- | --- | --- | --- | --- | --- |
| AW | 1,477 | 708 | 36.67 ± 11.85 | 1.50–94.33 | 50.2 | 49.8 |
| S-AW | 1,477 | 708 | 13.13 ± 3.92 | 0.80–32.33 | 46.6 | 53.4 |
| S-AW (REM) | 1,477 | 708 | 4.05 ± 2.04 | 0.00–13.00 | 39.7 | 60.3 |
| S-AW (REM) (Half1) | 1,477 | 708 | 1.18 ± 0.80 | 0.00–4.67 | 33.6 | 66.4 |
| S-AW (REM) (Half2) | 1,477 | 708 | 2.85 ± 1.52 | 0.00–8.67 | 35.6 | 64.4 |
| S-AW (NREM) | 1,477 | 708 | 7.90 ± 3.00 | 0.33–21.00 | 48.9 | 51.1 |
| S-AW (NREM) (Half1) | 1,477 | 708 | 3.95 ± 1.62 | 0.33–10.71 | 42.7 | 57.3 |
| S-AW (NREM) (Half2) | 1,477 | 708 | 3.92 ± 1.82 | 0.00–13.71 | 38.9 | 61.1 |
| M-AW | 1,477 | 708 | 13.53 ± 4.70 | 0.00–35.83 | 43.0 | 57.0 |
| M-AW (REM) | 1,477 | 708 | 4.32 ± 2.11 | 0.00–12.29 | 42.8 | 57.2 |
| M-AW (REM) (Half1) | 1,477 | 708 | 1.27 ± 0.84 | 0.00–5.00 | 34.8 | 65.2 |
| M-AW (REM) (Half2) | 1,477 | 708 | 3.04 ± 1.53 | 0.00–8.20 | 38.3 | 61.7 |
| M-AW (NREM) | 1,477 | 708 | 8.14 ± 3.27 | 0.00–21.33 | 43.6 | 56.4 |
| M-AW (NREM) (Half1) | 1,477 | 708 | 4.03 ± 1.78 | 0.00–11.75 | 40.4 | 59.6 |
| M-AW (NREM) (Half2) | 1,477 | 708 | 4.07 ± 1.86 | 0.00–14.33 | 39.2 | 60.8 |
| L-AW | 1,477 | 708 | 10.01 ± 5.94 | 0.00–31.00 | 50.2 | 49.8 |
| L-AW (REM) | 1,477 | 708 | 3.45 ± 2.51 | 0.00–13.83 | 53.6 | 46.4 |
| L-AW (REM) (Half1) | 1,477 | 708 | 0.98 ± 0.92 | 0.00–6.33 | 48.2 | 51.8 |
| L-AW (REM) (Half2) | 1,477 | 708 | 2.43 ± 1.80 | 0.00–10.83 | 47.4 | 52.6 |
| L-AW (NREM) | 1,477 | 708 | 5.31 ± 3.35 | 0.00–24.50 | 46.0 | 54.0 |
| L-AW (NREM) (Half1) | 1,477 | 708 | 2.85 ± 1.79 | 0.00–10.75 | 42.3 | 57.7 |
| L-AW (NREM) (Half2) | 1,477 | 708 | 2.40 ± 1.83 | 0.00–16.67 | 44.1 | 55.9 |
| FREM% | 1,477 | 708 | 43.96 ± 27.50 | 0.00–100.00 | 30.0 | 70.0 |
| TST | 1,477 | 708 | 357.97 ± 54.23 | 204.50–536.38 | 40.8 | 59.2 |

ICC and variance components estimated from unconditional random-intercept models: $Y \sim 1 + (1|\text{Participant})$ . TST = total sleep time (min); AW = objective awakening count; S-AW = short awakening count; M-AW = medium awakening count; L-AW = long awakening count; FREM% = percentage of final awakenings preceded by REM (%; 0–100); REM/NREM = sleep stage; Half1/Half2 = first/second half of the night.

**Table 4.** descriptive statistics of exploratory predictors + Other sleep variables in Dataset 1.

| Variable | N (obs) | N (ID) | Mean ± SD | Range | Between % | Within % |
| --- | --- | --- | --- | --- | --- | --- |
| WASO | 1,477 | 708 | 64.03 ± 34.48 | 0.83–236.67 | 44.2 | 55.8 |
| TIB | 1,477 | 708 | 457.24 ± 48.71 | 275.38–659.08 | 53.8 | 46.2 |
| Subj-AW | 1,477 | 708 | 1.22 ± 1.03 | 0.00–3.00 | 39.7 | 60.3 |
| Subj-TST | 1,438 | 701 | 2.14 ± 0.80 | 0.00–3.00 | 51.7 | 48.3 |
| REM | 1,477 | 708 | 85.72 ± 34.67 | 0.00–314.00 | 51.3 | 48.7 |
| REM (Half1) | 1,477 | 708 | 27.53 ± 13.80 | 0.00–142.83 | 48.1 | 51.9 |
| REM (Half2) | 1,477 | 708 | 58.19 ± 23.45 | 0.00–171.17 | 50.8 | 49.2 |
| N1 | 1,477 | 708 | 13.00 ± 16.68 | 0.50–207.33 | 3.2 | 96.8 |
| N1 (Half1) | 1,477 | 708 | 7.74 ± 7.76 | 0.00–90.21 | 12.3 | 87.7 |
| N1 (Half2) | 1,477 | 708 | 5.26 ± 9.92 | 0.00–124.33 | 3.8 | 96.2 |
| N2 | 1,477 | 708 | 116.66 ± 50.21 | 4.83–343.29 | 64.3 | 35.7 |
| N2 (Half1) | 1,477 | 708 | 45.70 ± 24.74 | 1.58–165.57 | 62.3 | 37.7 |
| N2 (Half2) | 1,477 | 708 | 70.97 ± 28.15 | 3.10–177.71 | 62.8 | 37.2 |
| N3 | 1,477 | 708 | 142.59 ± 44.83 | 15.83–284.07 | 65.5 | 34.5 |
| N3 (Half1) | 1,477 | 708 | 96.84 ± 26.53 | 12.17–181.07 | 57.5 | 42.5 |
| N3 (Half2) | 1,477 | 708 | 45.75 ± 22.82 | 0.17–127.38 | 63.2 | 36.8 |
| COFF | 1,413 | 688 | 0.25 ± 0.20 | 0.00–1.00 | 78.5 | 21.5 |
| ALCO | 1,413 | 688 | 0.06 ± 0.08 | 0.00–0.50 | 56.4 | 43.6 |
| CANN | 1,413 | 688 | 0.01 ± 0.03 | 0.00–0.49 | 76.7 | 23.3 |
| Anx-State | 1,355 | 658 | 33.30 ± 8.73 | 20.00–69.00 | 64.9 | 35.1 |
| DEP | 1,357 | 658 | 4.42 ± 3.12 | 0.00–18.00 | 56.8 | 43.2 |
| AM-S | 1,433 | 693 | 0.72 ± 0.16 | 0.00–1.00 | 25.8 | 74.2 |
| AM-L | 1,433 | 693 | 0.67 ± 0.15 | 0.00–1.00 | 21.3 | 78.7 |
| VWM | 1,032 | 462 | 7.72 ± 1.92 | 1.00–14.00 | 100.0 | 0.0 |
| AWM | 1,030 | 459 | 8.70 ± 2.42 | 0.00–14.00 | 100.0 | 0.0 |

ICC and variance components estimated from unconditional random-intercept models: $Y \sim 1 + (1|\text{Participant})$ ; WASO = wake after sleep onset (min); Subj-TST = subjective sleep duration (min); Subj-AW = subjective awakening count; REM = REM duration (min); N1 = N1 duration (min); N2 = N2 duration (min); N3 = N3 duration (min); COFF = proportion of beeps with coffee intake; ALCO = proportion of beeps with alcohol intake; CANN = proportion of beeps with cannabis intake; DEP = depression score; Anx-State = state anxiety score; AM-S = associative memory (short) accuracy; AM-L = associative memory (long) accuracy; VWM = visual working memory (forward); AWM = auditory working memory (forward). REM/NREM = sleep stage; Half1/Half2 = first/second half of the night.

**Table 5.** Descriptive statistics of main predictors in Dataset 2+3.

| Variable | N (obs) | N (ID) | Mean ± SD | Range | Between % | Within % |
| --- | --- | --- | --- | --- | --- | --- |
| AW | 2,896 | 124 | 25.76 ± 8.60 | 3.00–65.00 | 36.5 | 63.5 |
| S-AW | 2,896 | 124 | 12.22 ± 5.63 | 0.00–35.00 | 35.5 | 64.5 |
| S-AW (REM) | 2,896 | 124 | 3.32 ± 2.71 | 0.00–17.00 | 28.4 | 71.6 |
| S-AW (REM) (Half1) | 2,896 | 124 | 0.90 ± 1.23 | 0.00–9.00 | 15.4 | 84.6 |
| S-AW (REM) (Half2) | 2,896 | 124 | 2.41 ± 2.16 | 0.00–13.00 | 23.6 | 76.4 |
| S-AW (NREM) | 2,896 | 124 | 8.90 ± 4.59 | 0.00–31.00 | 35.9 | 64.1 |
| S-AW (NREM) (Half1) | 2,896 | 124 | 4.15 ± 2.72 | 0.00–18.00 | 24.7 | 75.3 |
| S-AW (NREM) (Half2) | 2,896 | 124 | 4.73 ± 2.84 | 0.00–18.00 | 26.4 | 73.6 |
| M-AW | 2,896 | 124 | 7.54 ± 3.53 | 0.00–23.00 | 20.7 | 79.3 |
| M-AW (REM) | 2,896 | 124 | 0.86 ± 1.02 | 0.00–7.00 | 13.4 | 86.6 |
| M-AW (REM) (Half1) | 2,896 | 124 | 0.24 ± 0.49 | 0.00–3.00 | 4.7 | 95.3 |
| M-AW (REM) (Half2) | 2,896 | 124 | 0.62 ± 0.84 | 0.00–7.00 | 9.8 | 90.2 |
| M-AW (NREM) | 2,896 | 124 | 6.67 ± 3.33 | 0.00–21.00 | 20.0 | 80.0 |
| M-AW (NREM) (Half1) | 2,896 | 124 | 3.08 ± 2.08 | 0.00–14.00 | 13.8 | 86.2 |
| M-AW (NREM) (Half2) | 2,896 | 124 | 3.57 ± 2.29 | 0.00–14.00 | 13.6 | 86.4 |
| L-AW | 2,896 | 124 | 6.01 ± 2.88 | 0.00–23.00 | 25.0 | 75.0 |
| L-AW (REM) | 2,896 | 124 | 0.81 ± 0.92 | 0.00–5.00 | 9.5 | 90.5 |
| L-AW (REM) (Half1) | 2,896 | 124 | 0.23 ± 0.48 | 0.00–3.00 | 5.0 | 95.0 |
| L-AW (REM) (Half2) | 2,896 | 124 | 0.57 ± 0.76 | 0.00–5.00 | 6.9 | 93.1 |
| L-AW (NREM) | 2,896 | 124 | 5.20 ± 2.72 | 0.00–21.00 | 24.3 | 75.7 |
| L-AW (NREM) (Half1) | 2,896 | 124 | 2.45 ± 1.95 | 0.00–12.00 | 20.7 | 79.3 |
| L-AW (NREM) (Half2) | 2,896 | 124 | 2.69 ± 1.89 | 0.00–20.00 | 12.6 | 87.4 |
| TST | 2,896 | 124 | 415.26 ± 81.61 | 155.00–900.00 | 20.5 | 79.5 |

ICC and variance components estimated from unconditional random-intercept models: $Y \sim 1 + (1|\text{Participant})$ . TST = total sleep time (min); AW = objective awakening count (count); S-AW = short awakenings (count); M-AW = medium awakenings (count); L-AW = long awakenings (count). REM/NREM = sleep stage; Half1/Half2 = first/second half of the night.

**Table 6.**
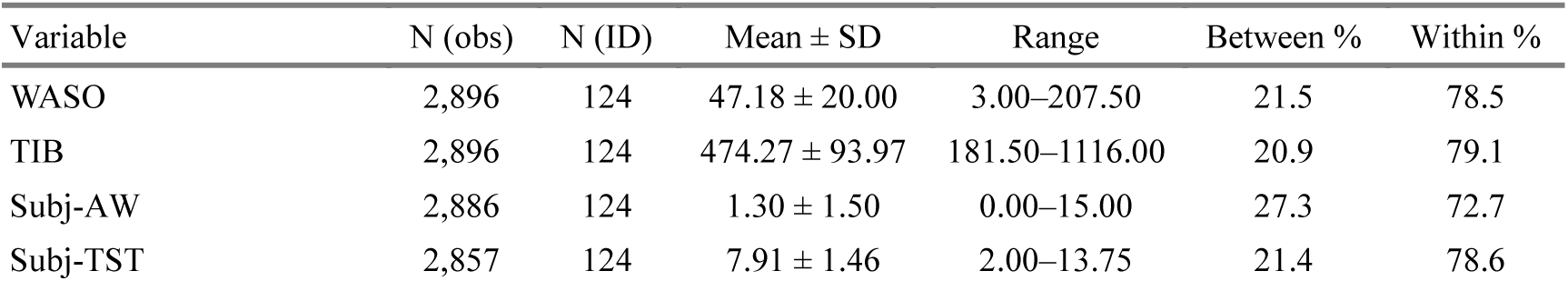
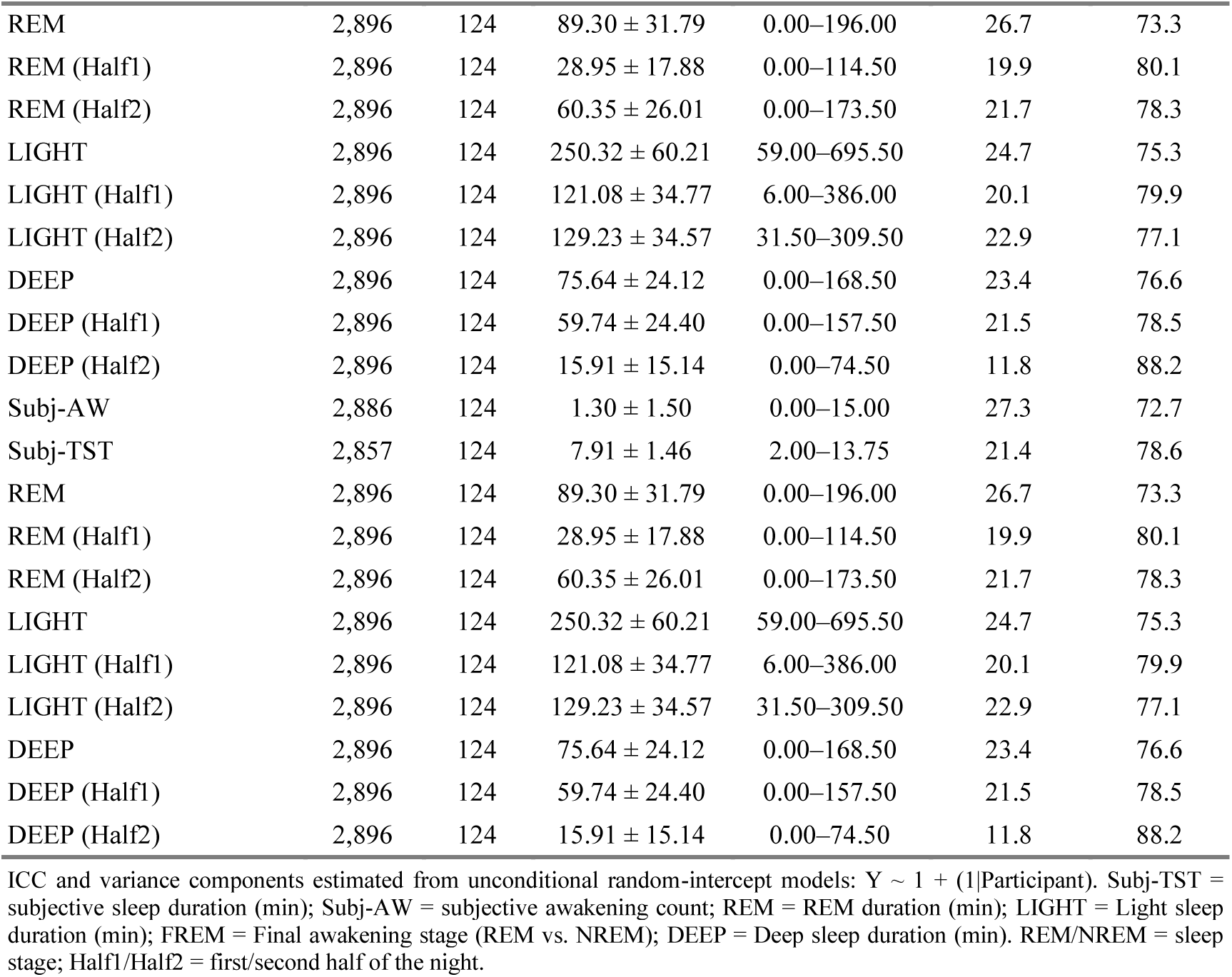
Descriptive statistics of Exploratory predictors + Other sleep variables in Dataset 2+3.

### Main and Exploratory Analyses Results

#### Models Predicting Dream Recall Frequency

##### Main Models

**Table 7.**
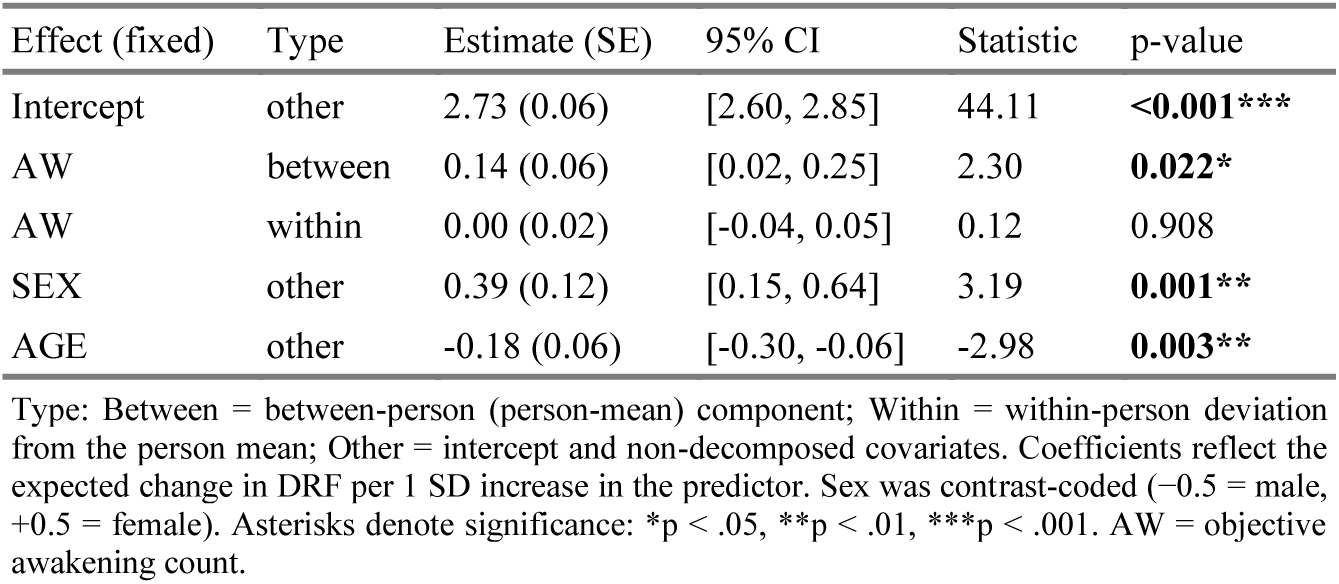
Model 1 (LME results for DRF)

**Table 8.** Model 2 (LME results for DRF)

| Effect (fixed) | Type | Estimate (SE) | 95% CI | Statistic | p-value |
| --- | --- | --- | --- | --- | --- |
| Intercept | other | 2.72 (0.06) | [2.60, 2.85] | 44.19 | **<0.001\*\*\*** |
| S-AW | between | 0.11 (0.08) | [-0.05, 0.26] | 1.36 | 0.175 |
| M-AW | between | -0.11 (0.10) | [-0.30, 0.08] | -1.18 | 0.238 |
| L-AW | between | 0.20 (0.08) | [0.04, 0.35] | 2.54 | **0.011\*** |
| S-AW | within | -0.01 (0.03) | [-0.06, 0.05] | -0.25 | 0.806 |
| M-AW | within | 0.07 (0.03) | [0.00, 0.13] | 2.12 | **0.035\*!** |
| L-AW | within | -0.06 (0.03) | [-0.12, -0.01] | -2.22 | **0.027\*** |
| SEX | other | 0.39 (0.12) | [0.15, 0.63] | 3.17 | **0.002\*\*** |
| AGE | other | -0.18 (0.06) | [-0.30, -0.06] | -2.99 | **0.003\*\*** |

Type: Between = between-person (person-mean) component; Within = within-person deviation from the person mean; Other = intercept and non-decomposed covariates. Coefficients reflect the expected change in DRF per 1 SD increase in the predictor. Sex was contrast-coded (-0.5 = male, +0.5 = female). Asterisks denote significance: \*p < .05, \*\*p < .01, \*\*\*p < .001. ! after the asterisks marks the effects that are not significant in the sensitivity analysis. S-AW = short awakening count; M-AW = medium awakening count; L-AW = long awakening count; REM/NREM = sleep stage; Half1/Half2 = first/second half of the night.

**Table 9.**
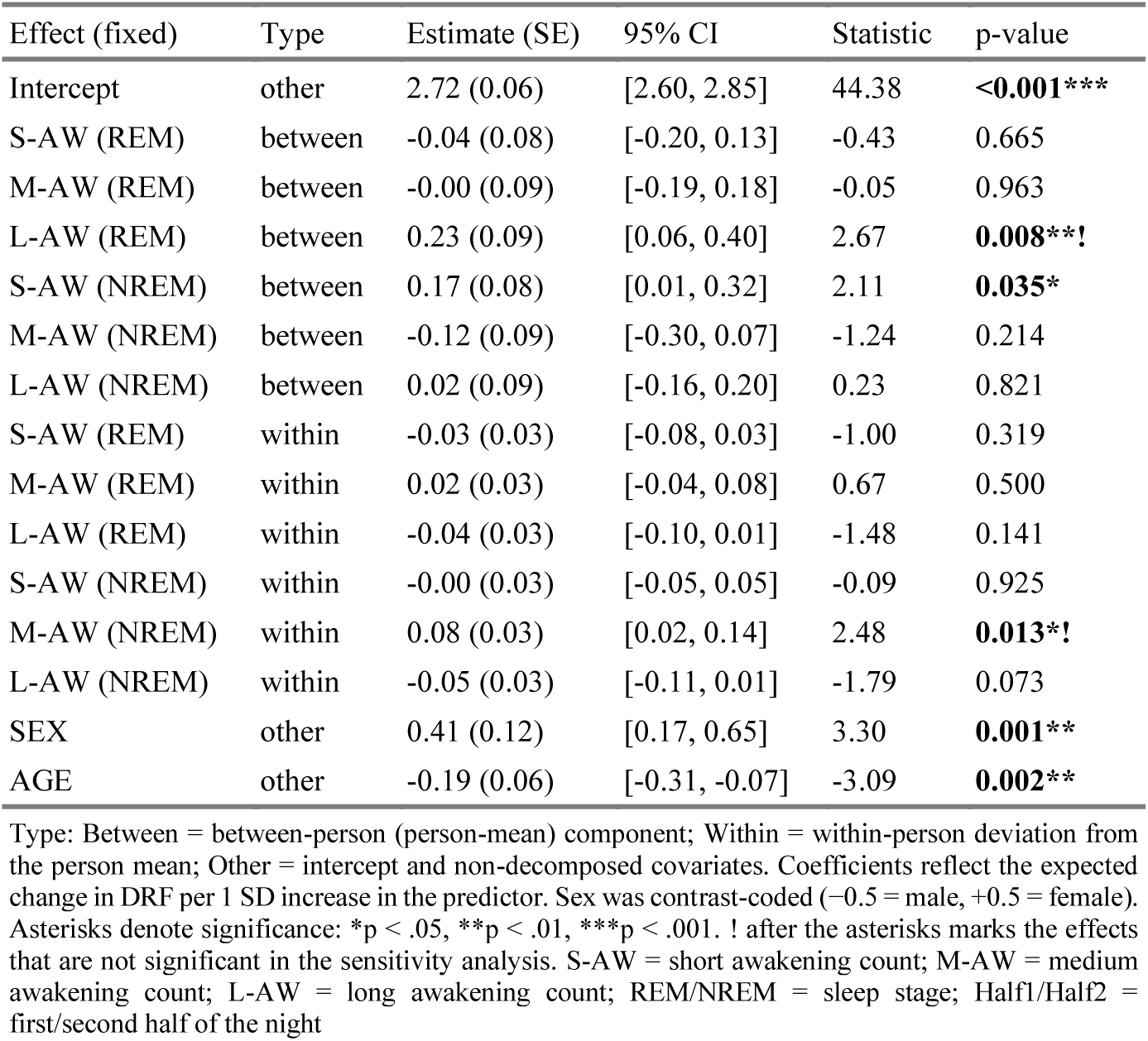
Model 3 (LME results for DRF)

| Effect (fixed) | Type | Estimate (SE) | 95% CI | Statistic | p-value |
| --- | --- | --- | --- | --- | --- |
| Intercept | other | 2.72 (0.06) | [2.60, 2.85] | 44.38 | **<0.001\*\*\*** |
| S-AW (REM) | between | -0.04 (0.08) | [-0.20, 0.13] | -0.43 | 0.665 |
| M-AW (REM) | between | -0.00 (0.09) | [-0.19, 0.18] | -0.05 | 0.963 |
| L-AW (REM) | between | 0.23 (0.09) | [0.06, 0.40] | 2.67 | **0.008\*\*!** |
| S-AW (NREM) | between | 0.17 (0.08) | [0.01, 0.32] | 2.11 | **0.035\*** |
| M-AW (NREM) | between | -0.12 (0.09) | [-0.30, 0.07] | -1.24 | 0.214 |
| L-AW (NREM) | between | 0.02 (0.09) | [-0.16, 0.20] | 0.23 | 0.821 |
| S-AW (REM) | within | -0.03 (0.03) | [-0.08, 0.03] | -1.00 | 0.319 |
| M-AW (REM) | within | 0.02 (0.03) | [-0.04, 0.08] | 0.67 | 0.500 |
| L-AW (REM) | within | -0.04 (0.03) | [-0.10, 0.01] | -1.48 | 0.141 |
| S-AW (NREM) | within | -0.00 (0.03) | [-0.05, 0.05] | -0.09 | 0.925 |
| M-AW (NREM) | within | 0.08 (0.03) | [0.02, 0.14] | 2.48 | **0.013\*!** |
| L-AW (NREM) | within | -0.05 (0.03) | [-0.11, 0.01] | -1.79 | 0.073 |
| SEX | other | 0.41 (0.12) | [0.17, 0.65] | 3.30 | **0.001\*\*** |
| AGE | other | -0.19 (0.06) | [-0.31, -0.07] | -3.09 | **0.002\*\*** |

Type: Between = between-person (person-mean) component; Within = within-person deviation from the person mean; Other = intercept and non-decomposed covariates. Coefficients reflect the expected change in DRF per 1 SD increase in the predictor. Sex was contrast-coded (-0.5 = male, +0.5 = female). Asterisks denote significance: \*p < .05, \*\*p < .01, \*\*\*p < .001. ! after the asterisks marks the effects that are not significant in the sensitivity analysis. S-AW = short awakening count; M-AW = medium awakening count; L-AW = long awakening count; REM/NREM = sleep stage; Half1/Half2 = first/second half of the night

**Table 10.** Model 4 (LME results for DRF)

| Effect (fixed) | Type | Estimate (SE) | 95% CI | Statistic | p-value |
| --- | --- | --- | --- | --- | --- |
| Intercept | other | 2.72 (0.06) | [2.60, 2.84] | 44.55 | **<0.001\*\*\*** |
| S-AW (REM) | between | -0.10 (0.09) | [-0.27, 0.07] | -1.21 | 0.229 |
| M-AW (REM) | between | -0.03 (0.09) | [-0.22, 0.15] | -0.33 | 0.740 |
| L-AW (REM) | between | 0.27 (0.09) | [0.09, 0.44] | 2.97 | **0.003\*\*** |
| S-AW (NREM) | between | 0.15 (0.08) | [-0.02, 0.31] | 1.77 | 0.077 |
| M-AW (NREM) | between | -0.10 (0.09) | [-0.29, 0.09] | -1.06 | 0.291 |
| L-AW (NREM) | between | 0.07 (0.09) | [-0.11, 0.25] | 0.73 | 0.469 |
| TST | between | 0.14 (0.07) | [-0.00, 0.29] | 1.93 | 0.053 |
| FREM% | between | 0.11 (0.07) | [-0.03, 0.24] | 1.59 | 0.113 |
| S-AW (REM) | within | -0.03 (0.03) | [-0.09, 0.02] | -1.18 | 0.239 |
| M-AW (REM) | within | 0.02 (0.03) | [-0.04, 0.08] | 0.60 | 0.547 |
| L-AW (REM) | within | -0.04 (0.03) | [-0.09, 0.02] | -1.35 | 0.178 |
| S-AW (NREM) | within | -0.00 (0.03) | [-0.06, 0.05] | -0.17 | 0.864 |
| M-AW (NREM) | within | 0.08 (0.03) | [0.02, 0.14] | 2.51 | **0.012\*!** |
| L-AW (NREM) | within | -0.05 (0.03) | [-0.11, 0.02] | -1.45 | 0.146 |
| TST | within | 0.02 (0.03) | [-0.04, 0.08] | 0.60 | 0.549 |
| FREM% | within | 0.01 (0.02) | [-0.04, 0.06] | 0.39 | 0.698 |
| SEX | other | 0.44 (0.12) | [0.19, 0.68] | 3.51 | **<0.001\*\*\*** |
| AGE | other | -0.20 (0.06) | [-0.32, -0.08] | -3.36 | **<0.001\*\*\*** |

Type: Between = between-person (person-mean) component; Within = within-person deviation from the person mean; Other = intercept and non-decomposed covariates. Coefficients reflect the expected change in DRF per 1 SD increase in the predictor. Sex was contrast-coded (-0.5 = male, +0.5 = female). Asterisks denote significance: \*p < .05, \*\*p < .01, \*\*\*p < .001. ! after the asterisks marks the effects that are not significant in the sensitivity analysis. TST = total sleep time; S-AW = short awakening count; M-AW = medium awakening count; L-AW = long awakening count; FREM% = percentage of final awakenings preceded by REM; REM/NREM = sleep stage; Half1/Half2 = first/second half of the night.

##### Exploratory Models

**Table 11.** Model E1 (LME results for DRF)

| Effect (fixed) | Type | Estimate (SE) | 95% CI | Statistic | p-value |
| --- | --- | --- | --- | --- | --- |
| Intercept | other | 2.72 (0.06) | [2.60, 2.84] | 44.38 | **<0.001\*\*\*** |
| S-AW (Half2) (REM) | between | -0.02 (0.08) | [-0.18, 0.14] | -0.21 | 0.832 |
| M-AW (Half2) (REM) | between | -0.05 (0.09) | [-0.23, 0.12] | -0.60 | 0.548 |
| L-AW (Half2) (REM) | between | 0.22 (0.08) | [0.05, 0.38] | 2.62 | **0.009\*\*** |
| S-AW (Half2) (NREM) | between | 0.06 (0.08) | [-0.09, 0.21] | 0.84 | 0.401 |
| M-AW (Half2) (NREM) | between | -0.02 (0.09) | [-0.19, 0.15] | -0.20 | 0.842 |
| L-AW (Half2) (NREM) | between | 0.07 (0.09) | [-0.10, 0.24] | 0.77 | 0.443 |
| TST | between | 0.13 (0.07) | [-0.01, 0.28] | 1.79 | 0.074 |
| FREM% | between | 0.10 (0.07) | [-0.04, 0.23] | 1.44 | 0.150 |
| S-AW (Half2) (REM) | within | -0.04 (0.03) | [-0.10, 0.02] | -1.40 | 0.161 |
| M-AW (Half2) (REM) | within | 0.01 (0.03) | [-0.05, 0.07] | 0.34 | 0.736 |
| L-AW (Half2) (REM) | within | -0.03 (0.03) | [-0.08, 0.02] | -1.06 | 0.287 |
| S-AW (Half2) (NREM) | within | 0.00 (0.03) | [-0.05, 0.05] | 0.07 | 0.947 |
| M-AW (Half2) (NREM) | within | 0.06 (0.03) | [0.01, 0.12] | 2.24 | **0.026\*!** |
| L-AW (Half2) (NREM) | within | -0.04 (0.03) | [-0.09, 0.02] | -1.20 | 0.230 |
| TST | within | 0.03 (0.03) | [-0.03, 0.08] | 0.89 | 0.373 |
| FREM% | within | 0.01 (0.02) | [-0.04, 0.06] | 0.36 | 0.722 |
| SEX | other | 0.45 (0.13) | [0.20, 0.69] | 3.58 | **<0.001\*\*\*** |
| AGE | other | -0.20 (0.06) | [-0.32, -0.08] | -3.23 | **0.001\*\*** |

Type: Between = between-person (person-mean) component; Within = within-person deviation from the person mean; Other = intercept and non-decomposed covariates. Coefficients reflect the expected change in DRF per 1 SD increase in the predictor. Sex was contrast-coded (-0.5 = male, +0.5 = female). Asterisks denote significance: \* $p < .05$ , \*\* $p < .01$ , \*\*\* $p < .001$ . ! after the asterisks marks the effects that are not significant in the sensitivity analysis. TST = total sleep time; S-AW = short awakening count; M-AW = medium awakening count; L-AW = long awakening count; FREM% = percentage of final awakenings preceded by REM; REM/NREM = sleep stage; Half1/Half2 = first/second half of the night.

**Table 12.** Model E2 (LME results for DRF)

| Effect (fixed) | Type | Estimate (SE) | 95% CI | Statistic | p-value |
| --- | --- | --- | --- | --- | --- |
| Intercept | other | 2.71 (0.06) | [2.59, 2.83] | 44.24 | **<0.001\*\*\*** |
| Subj-AW | between | 0.33 (0.06) | [0.21, 0.46] | 5.31 | **<0.001\*\*\*** |
| Subj-TST | between | 0.15 (0.06) | [0.03, 0.28] | 2.38 | **0.018\*** |
| Subj-AW | within | -0.02 (0.02) | [-0.07, 0.03] | -0.74 | 0.458 |
| Subj-TST | within | 0.02 (0.02) | [-0.03, 0.07] | 0.84 | 0.403 |
| SEX | other | 0.39 (0.12) | [0.14, 0.63] | 3.14 | **0.002\*\*** |
| AGE | other | -0.21 (0.06) | [-0.33, -0.09] | -3.50 | **<0.001\*\*\*** |

Type: Between = between-person (person-mean) component; Within = within-person deviation from the person mean; Other = intercept and non-decomposed covariates. Coefficients reflect the expected change in DRF per 1 SD increase in the predictor. Sex was contrast-coded (-0.5 = male, +0.5 = female). Asterisks denote significance: \* $p < .05$ , \*\* $p < .01$ , \*\*\* $p < .001$ . Abbreviations: Subj-TST = subjective sleep duration; Subj-AW = subjective awakening count.

**Table 13.** Model E3 (LME results for DRF)

| Effect (fixed) | Type | Estimate (SE) | 95% CI | Statistic | p-value |
| --- | --- | --- | --- | --- | --- |
| Intercept | other | 2.71 (0.08) | [2.56, 2.86] | 35.36 | **<0.001\*\*\*** |
| Subj-AW | between | 0.33 (0.08) | [0.16, 0.50] | 3.88 | **<0.001\*\*\*** |
| Subj-TST | between | 0.02 (0.08) | [-0.14, 0.18] | 0.23 | 0.817 |
| ALCO | between | 0.12 (0.09) | [-0.05, 0.28] | 1.36 | 0.174 |
| COFF | between | -0.12 (0.09) | [-0.29, 0.05] | -1.39 | 0.167 |
| CANN | between | 0.19 (0.07) | [0.05, 0.33] | 2.68 | **0.008\*\*** |
| DEP | between | -0.06 (0.11) | [-0.28, 0.16] | -0.55 | 0.580 |
| Anx-State | between | 0.02 (0.11) | [-0.19, 0.24] | 0.23 | 0.817 |
| AM-L | between | 0.02 (0.08) | [-0.14, 0.18] | 0.21 | 0.831 |
| AM-S | between | 0.03 (0.09) | [-0.14, 0.20] | 0.32 | 0.749 |
| Subj-AW | within | -0.00 (0.03) | [-0.07, 0.06] | -0.13 | 0.893 |
| Subj-TST | within | 0.02 (0.03) | [-0.04, 0.08] | 0.74 | 0.458 |
| ALCO | within | 0.06 (0.03) | [-0.00, 0.12] | 1.86 | 0.064 |
| COFF | within | -0.00 (0.03) | [-0.06, 0.06] | -0.07 | 0.945 |
| CANN | within | -0.01 (0.04) | [-0.09, 0.06] | -0.30 | 0.766 |
| DEP | within | -0.04 (0.03) | [-0.11, 0.02] | -1.37 | 0.170 |
| Anx-State | within | 0.03 (0.03) | [-0.04, 0.09] | 0.81 | 0.421 |
| AM-L | within | -0.01 (0.03) | [-0.07, 0.05] | -0.41 | 0.683 |
| AM-S | within | -0.01 (0.03) | [-0.07, 0.04] | -0.50 | 0.616 |
| VWM | other | 0.07 (0.08) | [-0.09, 0.23] | 0.88 | 0.381 |
| AWM | other | 0.05 (0.08) | [-0.10, 0.20] | 0.64 | 0.524 |
| SEX | other | 0.20 (0.16) | [-0.12, 0.52] | 1.25 | 0.213 |
| AGE | other | -0.21 (0.08) | [-0.36, -0.06] | -2.76 | **0.006\*\*** |

Type: Between = between-person (person-mean) component; Within = within-person deviation from the person mean; Other = intercept and non-decomposed covariates. Coefficients reflect the expected change in DRF per 1 SD increase in the predictor. Sex was contrast-coded (-0.5 = male, +0.5 = female). Asterisks denote significance: \*p < .05, \*\*p < .01, \*\*\*p < .001. Abbreviations: Subj-TST = subjective sleep duration; Subj-AW = subjective awakening count; COFF = proportion of beeps with coffee intake; ALCO = proportion of beeps with alcohol intake; CANN = proportion of beeps with cannabis intake; Anx-State = state anxiety score; DEP = depression score; AM-L = associative memory (long) accuracy; AM-S = associative memory (short) accuracy; VWM = visual working memory (forward); AWM = auditory working memory (forward).

##### Models Summaries

**Table 14.**
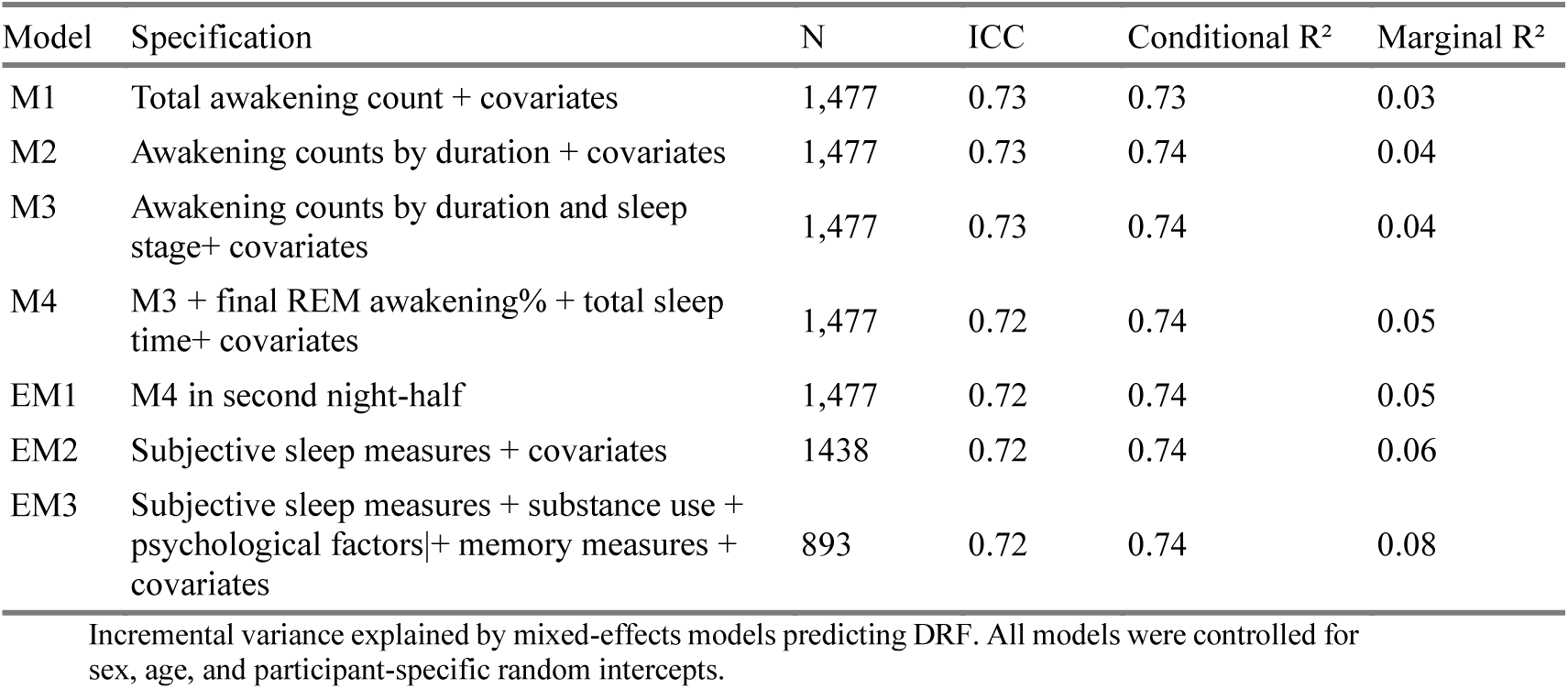
Model summaries for DRF.

#### Models Predicting Daily Dream Recall

##### Main Models

**Table 15.**
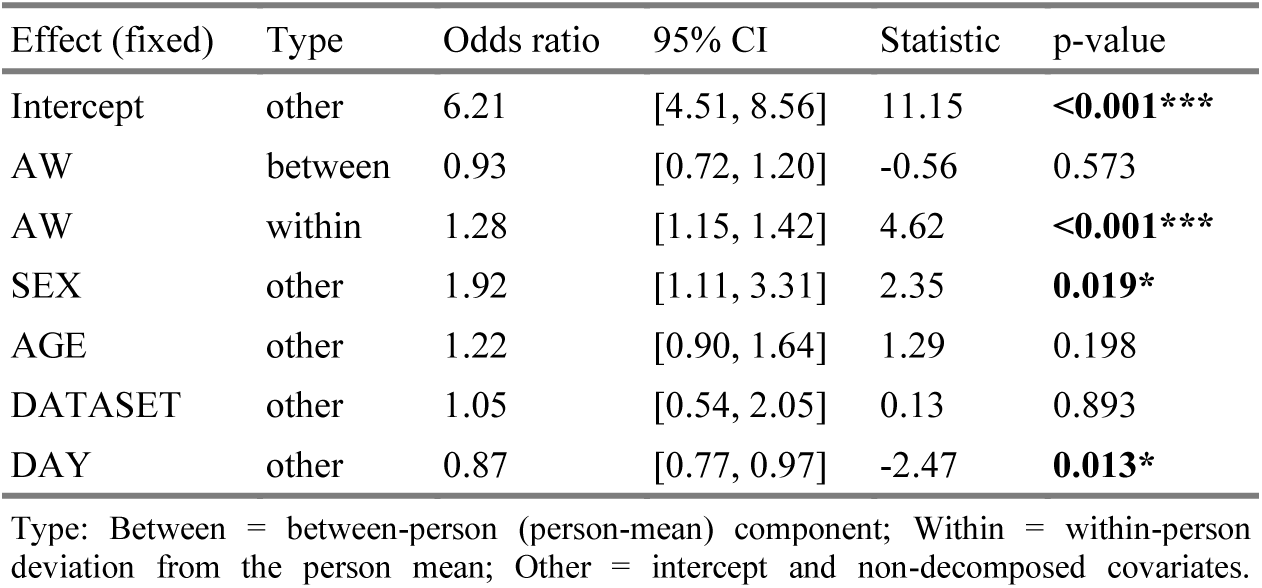
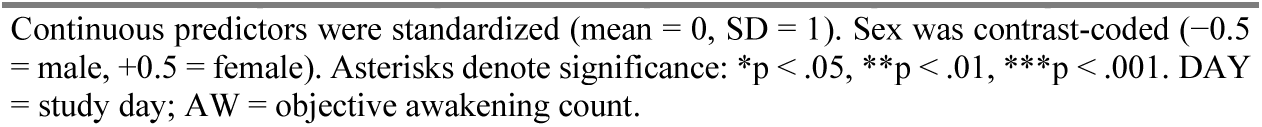
Model 1 (GLMM results for WD+CD vs. NE)

| Effect (fixed) | Type | Odds ratio | 95% CI | Statistic | p-value |
| --- | --- | --- | --- | --- | --- |
| Intercept | other | 6.21 | [4.51, 8.56] | 11.15 | **<0.001\*\*\*** |
| AW | between | 0.93 | [0.72, 1.20] | -0.56 | 0.573 |
| AW | within | 1.28 | [1.15, 1.42] | 4.62 | **<0.001\*\*\*** |
| SEX | other | 1.92 | [1.11, 3.31] | 2.35 | **0.019\*** |
| AGE | other | 1.22 | [0.90, 1.64] | 1.29 | 0.198 |
| DATASET | other | 1.05 | [0.54, 2.05] | 0.13 | 0.893 |
| DAY | other | 0.87 | [0.77, 0.97] | -2.47 | **0.013\*** |
| Continuous predictors were standardized (mean = 0, SD = 1). Sex was contrast-coded (-0.5 = male, +0.5 = female). Asterisks denote significance: *p < .05, **p < .01, ***p < .001. DAY = study day; AW = objective awakening count. |  |  |  |  |  |

**Table 16.** Model 2 (GLMM results for WD+CD vs. NE)

| Effect (fixed) | Type | Odds ratio | 95% CI | Statistic | p-value |
| --- | --- | --- | --- | --- | --- |
| Intercept | other | 6.34 | [4.60, 8.73] | 11.29 | **<0.001\*\*\*** |
| S-AW | between | 1.22 | [0.91, 1.64] | 1.31 | 0.190 |
| M-AW | between | 0.84 | [0.62, 1.13] | -1.14 | 0.253 |
| L-AW | between | 0.83 | [0.63, 1.09] | -1.38 | 0.169 |
| S-AW | within | 1.22 | [1.09, 1.36] | 3.57 | **<0.001\*\*\*** |
| M-AW | within | 1.08 | [0.97, 1.20] | 1.37 | 0.171 |
| L-AW | within | 1.09 | [0.98, 1.21] | 1.53 | 0.127 |
| SEX | other | 1.98 | [1.14, 3.42] | 2.43 | **0.015\*** |
| AGE | other | 1.19 | [0.88, 1.61] | 1.13 | 0.258 |
| DATASET | other | 0.97 | [0.50, 1.89] | -0.09 | 0.925 |
| DAY | other | 0.87 | [0.77, 0.97] | -2.45 | **0.014\*** |

Type: Between = between-person (person-mean) component; Within = within-person deviation from the person mean; Other = intercept and non-decomposed covariates. Continuous predictors were standardized (mean = 0, SD = 1) unless inherently coded. Sex was contrast-coded (-0.5 = male, +0.5 = female). Asterisks denote significance: \*p < .05, \*\*p < .01, \*\*\*p < .001. DAY = study day; S-AW = short awakening count; M-AW = medium awakening count; L-AW = long awakening count; REM/NREM = sleep stage.

**Table 17.** Model 3 (GLMM results for WD+CD vs. NE)

| Effect (fixed) | Type | Odds ratio | 95% CI | Statistic | p-value |
| --- | --- | --- | --- | --- | --- |
| Intercept | other | 6.07 | [4.43, 8.31] | 11.24 | **<0.001\*\*\*** |
| S-AW (REM) | between | 0.80 | [0.56, 1.14] | -1.25 | 0.210 |
| M-AW (REM) | between | 1.00 | [0.69, 1.45] | -0.01 | 0.994 |
| L-AW (REM) | between | 1.38 | [1.05, 1.80] | 2.34 | **0.019\*** |
| S-AW (NREM) | between | 1.46 | [1.08, 1.99] | 2.42 | **0.015\*** |
| M-AW (NREM) | between | 0.86 | [0.64, 1.17] | -0.95 | 0.344 |
| L-AW (NREM) | between | 0.68 | [0.51, 0.91] | -2.65 | **0.008\*\*** |
| S-AW (REM) | within | 1.23 | [1.10, 1.38] | 3.62 | **<0.001\*\*\*** |
| M-AW (REM) | within | 1.08 | [0.97, 1.20] | 1.47 | 0.142 |
| L-AW (REM) | within | 1.11 | [1.00, 1.24] | 1.92 | 0.055 |
| S-AW (NREM) | within | 1.09 | [0.97, 1.21] | 1.47 | 0.142 |
| M-AW (NREM) | within | 1.05 | [0.95, 1.17] | 0.95 | 0.340 |
| L-AW (NREM) | within | 1.07 | [0.96, 1.19] | 1.16 | 0.247 |
| SEX | other | 2.29 | [1.30, 4.06] | 2.85 | **0.004\*\*** |
| AGE | other | 1.21 | [0.90, 1.62] | 1.25 | 0.212 |
| DATASET | other | 1.11 | [0.57, 2.14] | 0.31 | 0.756 |
| DAY | other | 0.87 | [0.77, 0.98] | -2.34 | **0.019\*** |

| Effect (fixed) | Type | Odds ratio | 95% CI | Statistic | p-value |
| --- | --- | --- | --- | --- | --- |
| awakening count; M-AW = medium awakening count; L-AW = long awakening count; REM/NREM = sleep stage; Half1/Half2 = first/second half of the night. |  |  |  |  |  |

**Table 18.** Model 4 (GLMM results for WD+CD vs. NE)

| Effect (fixed) | Type | Odds ratio | 95% CI | Statistic | p-value |
| --- | --- | --- | --- | --- | --- |
| Intercept | other | 7.26 | [5.21, 10.12] | 11.70 | **<0.001\*\*\*** |
| S-AW (REM) | between | 0.77 | [0.53, 1.10] | -1.44 | 0.150 |
| M-AW (REM) | between | 1.03 | [0.70, 1.51] | 0.16 | 0.874 |
| L-AW (REM) | between | 1.33 | [1.01, 1.76] | 2.04 | **0.042\*** |
| S-AW (NREM) | between | 1.41 | [1.03, 1.94] | 2.14 | **0.032\*** |
| M-AW (NREM) | between | 0.81 | [0.58, 1.12] | -1.28 | 0.201 |
| L-AW (NREM) | between | 0.67 | [0.50, 0.89] | -2.75 | **0.006\*\*** |
| TST | between | 1.19 | [0.86, 1.66] | 1.06 | 0.289 |
| S-AW (REM) | within | 1.18 | [1.05, 1.33] | 2.69 | **0.007\*\*** |
| M-AW (REM) | within | 1.07 | [0.96, 1.19] | 1.17 | 0.241 |
| L-AW (REM) | within | 1.10 | [0.98, 1.23] | 1.62 | 0.105 |
| S-AW (NREM) | within | 1.06 | [0.94, 1.20] | 0.95 | 0.341 |
| M-AW (NREM) | within | 1.03 | [0.92, 1.16] | 0.48 | 0.631 |
| L-AW (NREM) | within | 1.03 | [0.92, 1.16] | 0.54 | 0.592 |
| TST | within | 1.11 | [0.95, 1.29] | 1.29 | 0.195 |
| PREM | other | 1.69 | [1.26, 2.27] | 3.48 | **<0.001\*\*\*** |
| SEX | other | 2.13 | [1.17, 3.87] | 2.48 | **0.013\*** |
| AGE | other | 1.18 | [0.88, 1.60] | 1.11 | 0.268 |
| DATASET | other | 1.17 | [0.60, 2.28] | 0.46 | 0.643 |
| DAY | other | 0.87 | [0.77, 0.98] | -2.29 | **0.022\*** |

Type: Between = between-person (person-mean) component; Within = within-person deviation from the person mean; Other = intercept and non-decomposed covariates. Continuous predictors were standardized (mean = 0, SD = 1). Sex was contrast-coded (-0.5 = male, +0.5 = female). Asterisks denote significance: \*p < .05, \*\*p < .01, \*\*\*p < .001. DATASET = Dataset vs. 2; DAY = study day; TST = total sleep time; S-AW = short awakening count; M-AW = medium awakening count; L-AW = long awakening count; PREM = final awakening stage (PREM); REM/NREM = sleep stage.

##### Exploratory Models

**Table 19.** Model E1 (GLMM results for WD+CD vs. NE)

| Effect (fixed) | Type | Odds ratio | 95% CI | Statistic | p-value |
| --- | --- | --- | --- | --- | --- |
| Intercept | other | 7.41 | [5.32, 10.32] | 11.84 | **<0.001\*\*\*** |
| S-AW (Half2) (REM) | between | 0.87 | [0.63, 1.21] | -0.81 | 0.418 |
| M-AW (Half2) (REM) | between | 0.82 | [0.58, 1.16] | -1.11 | 0.266 |
| L-AW (Half2) (REM) | between | 1.38 | [1.05, 1.81] | 2.34 | **0.019\*** |
| S-AW (Half2) (NREM) | between | 1.32 | [0.98, 1.78] | 1.82 | 0.068 |
| M-AW (Half2) (NREM) | between | 0.92 | [0.67, 1.27] | -0.49 | 0.626 |
| L-AW (Half2) (NREM) | between | 0.76 | [0.58, 1.00] | -1.95 | 0.051 |
| TST | between | 1.09 | [0.79, 1.50] | 0.50 | 0.614 |
| S-AW (Half2) (REM) | within | 1.15 | [1.02, 1.29] | 2.30 | **0.021\*** |
| M-AW (Half2) (REM) | within | 1.13 | [1.01, 1.26] | 2.18 | **0.029\*** |
| L-AW (Half2) (REM) | within | 1.07 | [0.96, 1.19] | 1.14 | 0.254 |
| S-AW (Half2) (NREM) | within | 1.08 | [0.96, 1.21] | 1.32 | 0.188 |
| M-AW (Half2) (NREM) | within | 1.03 | [0.92, 1.15] | 0.49 | 0.624 |
| L-AW (Half2) (NREM) | within | 1.10 | [0.98, 1.23] | 1.66 | 0.098 |
| TST | within | 1.11 | [0.98, 1.27] | 1.60 | 0.110 |
| PREM | other | 1.70 | [1.27, 2.27] | 3.57 | **<0.001\*\*\*** |
| SEX | other | 2.05 | [1.13, 3.73] | 2.36 | **0.018\*** |
| AGE | other | 1.14 | [0.84, 1.55] | 0.82 | 0.411 |
| DATASET | other | 1.06 | [0.54, 2.09] | 0.16 | 0.869 |
| DAY | other | 0.87 | [0.78, 0.98] | -2.35 | **0.019\*** |

Type: Between = between-person (person-mean) component; Within = within-person deviation from the person mean; Other = intercept and non-decomposed covariates. Continuous predictors were standardized (mean = 0, SD = 1). Sex was contrast-coded (-0.5 = male, +0.5 = female). Asterisks denote significance: \*p < .05, \*\*p < .01, \*\*\*p < .001. DAY = study day; TST = total sleep time; S-AW = short awakening count; M-AW = medium awakening count; L-AW = long awakening count; PREM = final awakening stage (PREM); REM/NREM = sleep stage; Half1/Half2 = first/second half of the night.

**Table 20.** Model E2 (GLMM results for WD+CD vs. NE)

| Effect (fixed) | Type | Odds ratio | 95% CI | Statistic | p-value |
| --- | --- | --- | --- | --- | --- |
| Intercept | other | 6.51 | [4.72, 8.99] | 11.40 | **<0.001\*\*\*** |
| Subj-AW | between | 1.38 | [1.04, 1.84] | 2.20 | **0.028\*** |
| Subj-TST | between | 1.02 | [0.77, 1.36] | 0.17 | 0.868 |
| Subj-AW | within | 1.30 | [1.15, 1.47] | 4.10 | **<0.001\*\*\*** |
| Subj-TST | within | 1.30 | [1.16, 1.44] | 4.73 | **<0.001\*\*\*** |
| SEX | other | 1.74 | [0.99, 3.07] | 1.93 | 0.054 |
| AGE | other | 1.13 | [0.84, 1.53] | 0.81 | 0.415 |
| DATASET | other | 1.14 | [0.58, 2.26] | 0.38 | 0.704 |
| DAY | other | 0.87 | [0.77, 0.98] | -2.30 | **0.021\*** |

Type: Between = between-person (person-mean) component; Within = within-person deviation from the person mean; Other = intercept and non-decomposed covariates. Continuous predictors were standardized (mean = 0, SD = 1). Sex was contrast-coded (-0.5 = male, +0.5 = female). Asterisks denote significance: \*p < .05, \*\*p < .01, \*\*\*p < .001. DAY = study day; Subj-TST = subjective sleep duration; Subj-AW = subjective awakening count;

##### Models Summaries

**Table 21.**
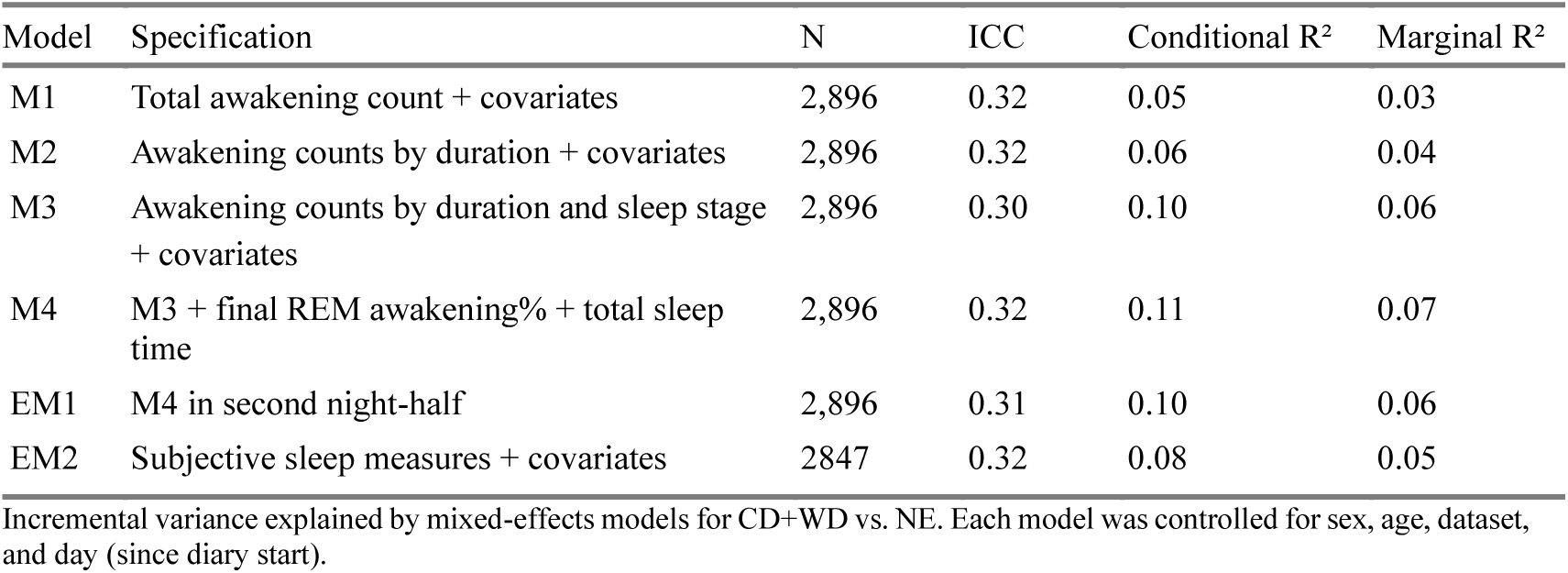
Model summaries for dream recall.

| Model | Specification | N | ICC | Conditional R<sup>2</sup> | Marginal R<sup>2</sup> |
| --- | --- | --- | --- | --- | --- |
| M1 | Total awakening count + covariates | 2,896 | 0.32 | 0.05 | 0.03 |
| M2 | Awakening counts by duration + covariates | 2,896 | 0.32 | 0.06 | 0.04 |
| M3 | Awakening counts by duration and sleep stage + covariates | 2,896 | 0.30 | 0.10 | 0.06 |
| M4 | M3 + final REM awakening% + total sleep time | 2,896 | 0.32 | 0.11 | 0.07 |
| EM1 | M4 in second night-half | 2,896 | 0.31 | 0.10 | 0.06 |
| EM2 | Subjective sleep measures + covariates | 2847 | 0.32 | 0.08 | 0.05 |

| Model | Specification | N | ICC | Conditional R<sup>2</sup> | Marginal R<sup>2</sup> |
| --- | --- | --- | --- | --- | --- |

#### Models Predicting Daily Content Recall

##### Main Models

**Table 22.**
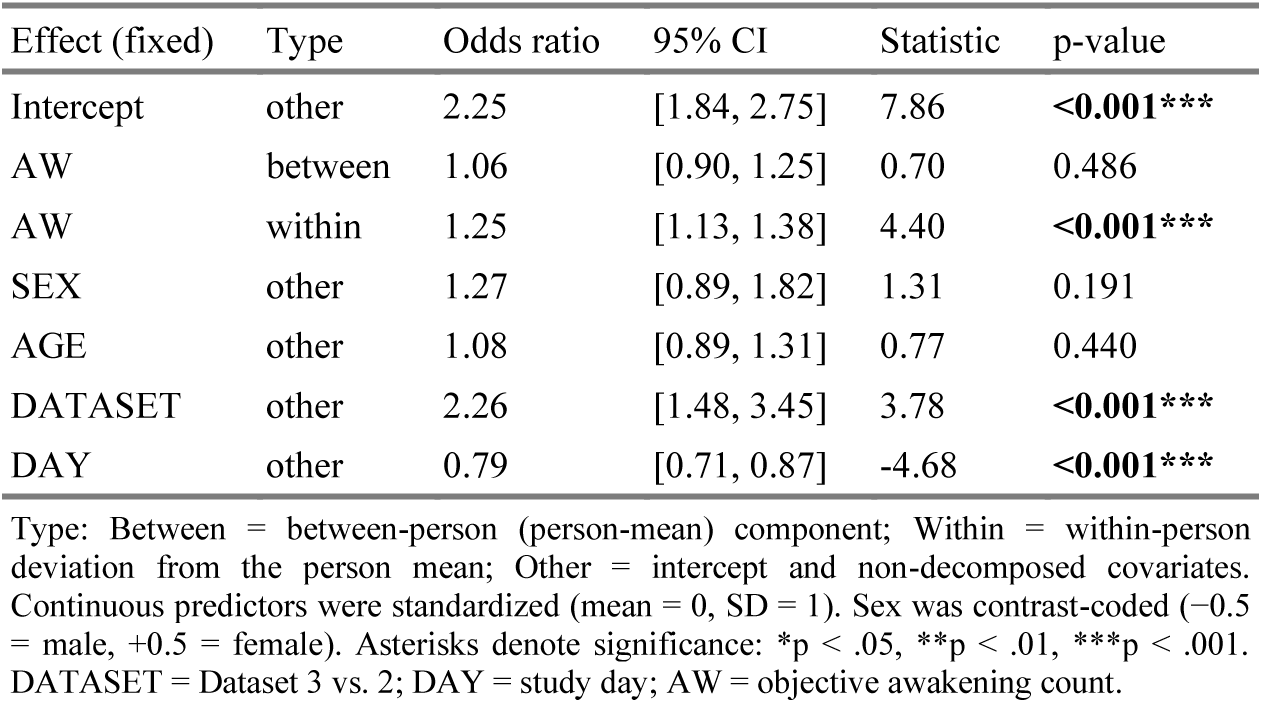
Model 1 (GLMM results for CD vs. WD)

**Table 23.** Model 2 (GLMM results for CD vs. WD)

| Effect (fixed) | Type | Odds ratio | 95% CI | Statistic | p-value |
| --- | --- | --- | --- | --- | --- |
| Intercept | other | 2.25 | [1.84, 2.75] | 7.93 | **<0.001\*\*\*** |
| S-AW | between | 1.12 | [0.93, 1.36] | 1.18 | 0.240 |
| M-AW | between | 0.94 | [0.77, 1.13] | -0.68 | 0.496 |
| L-AW | between | 1.01 | [0.84, 1.21] | 0.06 | 0.956 |
| S-AW | within | 1.14 | [1.04, 1.26] | 2.67 | **0.008\*\*** |
| M-AW | within | 1.10 | [0.99, 1.21] | 1.79 | 0.074 |
| L-AW | within | 1.12 | [1.02, 1.24] | 2.27 | **0.023\*** |
| SEX | other | 1.30 | [0.90, 1.86] | 1.42 | 0.157 |
| AGE | other | 1.07 | [0.89, 1.30] | 0.73 | 0.468 |
| DATASET | other | 2.20 | [1.44, 3.35] | 3.65 | **<0.001\*\*\*** |
| DAY | other | 0.79 | [0.71, 0.87] | -4.62 | **<0.001\*\*\*** |

Type: Between = between-person (person-mean) component; Within = within-person deviation from the person mean; Other = intercept and non-decomposed covariates. Continuous predictors were standardized (mean = 0, SD = 1). Sex was contrast-coded (-0.5 = male, +0.5 = female). Asterisks denote significance: \*p < .05, \*\*p < .01, \*\*\*p < .001. DATASET = Dataset 3 vs. 2; DAY = study day; S-AW = short awakening count; M-AW = medium awakening count; L-AW = long awakening count.

**Table 24.** Model 3 (GLMM results for CD vs. WD)

| Effect (fixed) | Type | Odds ratio | 95% CI | Statistic | p-value |
| --- | --- | --- | --- | --- | --- |
| Intercept | other | 2.27 | [1.87, 2.76] | 8.26 | **<0.001\*\*\*** |
| S-AW (REM) | between | 1.18 | [0.94, 1.50] | 1.42 | 0.157 |
| M-AW (REM) | between | 0.72 | [0.56, 0.92] | -2.59 | **0.009\*\*** |
| L-AW (REM) | between | 1.19 | [0.99, 1.42] | 1.87 | 0.061 |
| S-AW (NREM) | between | 1.09 | [0.90, 1.31] | 0.91 | 0.365 |
| M-AW (NREM) | between | 1.03 | [0.86, 1.25] | 0.33 | 0.739 |
| L-AW (NREM) | between | 0.99 | [0.82, 1.18] | -0.13 | 0.896 |
| S-AW (REM) | within | 1.01 | [0.91, 1.11] | 0.12 | 0.908 |
| M-AW (REM) | within | 1.08 | [0.98, 1.19] | 1.51 | 0.130 |
| L-AW (REM) | within | 1.14 | [1.04, 1.26] | 2.72 | **0.006\*\*** |
| S-AW (NREM) | within | 1.16 | [1.05, 1.28] | 2.92 | **0.003\*\*** |
| M-AW (NREM) | within | 1.08 | [0.98, 1.19] | 1.50 | 0.134 |
| L-AW (NREM) | within | 1.07 | [0.97, 1.18] | 1.40 | 0.160 |
| SEX | other | 1.19 | [0.82, 1.71] | 0.91 | 0.360 |
| AGE | other | 1.12 | [0.93, 1.35] | 1.17 | 0.244 |
| DATASET | other | 2.22 | [1.47, 3.34] | 3.82 | **<0.001\*\*\*** |
| DAY | other | 0.79 | [0.71, 0.87] | -4.60 | **<0.001\*\*\*** |

Type: Between = between-person (person-mean) component; Within = within-person deviation from the person mean; Other = intercept and non-decomposed covariates. Continuous predictors were standardized (mean = 0, SD = 1) unless inherently coded. Sex was contrast-coded (-0.5 = male, +0.5 = female). Asterisks denote significance: \* $p < .05$ , \*\* $p < .01$ , \*\*\* $p < .001$ . DATASET = Dataset 3 vs. 2; DAY = study day; S-AW = short awakening count; M-AW = medium awakening count; L-AW = long awakening count; REM/NREM = sleep stage.

**Table 25.**
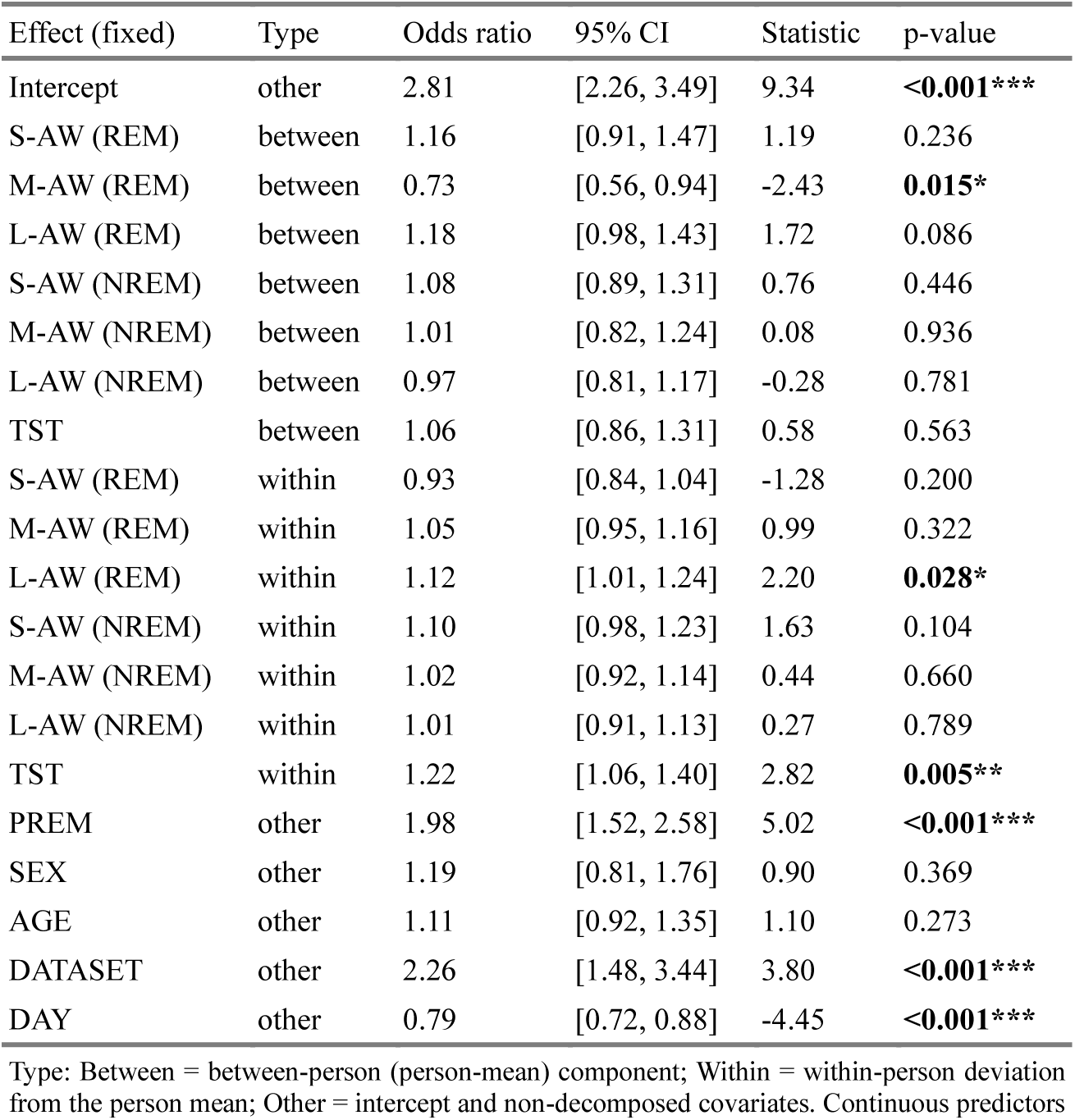
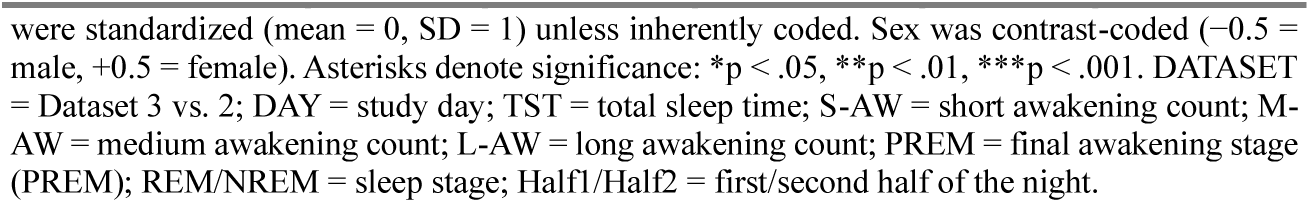
Model 4 (GLMM results for CD vs. WD)

| Effect (fixed) | Type | Odds ratio | 95% CI | Statistic | p-value |
| --- | --- | --- | --- | --- | --- |
| Intercept | other | 2.81 | [2.26, 3.49] | 9.34 | **<0.001\*\*\*** |
| S-AW (REM) | between | 1.16 | [0.91, 1.47] | 1.19 | 0.236 |
| M-AW (REM) | between | 0.73 | [0.56, 0.94] | -2.43 | **0.015\*** |
| L-AW (REM) | between | 1.18 | [0.98, 1.43] | 1.72 | 0.086 |
| S-AW (NREM) | between | 1.08 | [0.89, 1.31] | 0.76 | 0.446 |
| M-AW (NREM) | between | 1.01 | [0.82, 1.24] | 0.08 | 0.936 |
| L-AW (NREM) | between | 0.97 | [0.81, 1.17] | -0.28 | 0.781 |
| TST | between | 1.06 | [0.86, 1.31] | 0.58 | 0.563 |
| S-AW (REM) | within | 0.93 | [0.84, 1.04] | -1.28 | 0.200 |
| M-AW (REM) | within | 1.05 | [0.95, 1.16] | 0.99 | 0.322 |
| L-AW (REM) | within | 1.12 | [1.01, 1.24] | 2.20 | **0.028\*** |
| S-AW (NREM) | within | 1.10 | [0.98, 1.23] | 1.63 | 0.104 |
| M-AW (NREM) | within | 1.02 | [0.92, 1.14] | 0.44 | 0.660 |
| L-AW (NREM) | within | 1.01 | [0.91, 1.13] | 0.27 | 0.789 |
| TST | within | 1.22 | [1.06, 1.40] | 2.82 | **0.005\*\*** |
| PREM | other | 1.98 | [1.52, 2.58] | 5.02 | **<0.001\*\*\*** |
| SEX | other | 1.19 | [0.81, 1.76] | 0.90 | 0.369 |
| AGE | other | 1.11 | [0.92, 1.35] | 1.10 | 0.273 |
| DATASET | other | 2.26 | [1.48, 3.44] | 3.80 | **<0.001\*\*\*** |
| DAY | other | 0.79 | [0.72, 0.88] | -4.45 | **<0.001\*\*\*** |
| were standardized (mean = 0, SD = 1) unless inherently coded. Sex was contrast-coded (-0.5 = male, +0.5 = female). Asterisks denote significance: *p < .05, **p < .01, ***p < .001. DATASET = Dataset 3 vs. 2; DAY = study day; TST = total sleep time; S-AW = short awakening count; M-AW = medium awakening count; L-AW = long awakening count; PREM = final awakening stage (PREM); REM/NREM = sleep stage; Half1/Half2 = first/second half of the night. |  |  |  |  |  |

##### Exploratory Models

**Table 26.** Model E1 (GLMM results for CD vs. WD)

| Effect (fixed) | Type | Odds ratio | 95% CI | Statistic | p-value |
| --- | --- | --- | --- | --- | --- |
| Intercept | other | 2.82 | [2.27, 3.49] | 9.41 | **<0.001\*\*\*** |
| S-AW (Half2) (REM) | between | 1.06 | [0.85, 1.32] | 0.52 | 0.604 |
| M-AW (Half2) (REM) | between | 0.77 | [0.62, 0.97] | -2.23 | **0.026\*** |
| L-AW (Half2) (REM) | between | 1.14 | [0.95, 1.36] | 1.39 | 0.165 |
| S-AW (Half2) (NREM) | between | 1.15 | [0.95, 1.38] | 1.41 | 0.160 |
| M-AW (Half2) (NREM) | between | 0.98 | [0.80, 1.20] | -0.20 | 0.843 |
| L-AW (Half2) (NREM) | between | 0.99 | [0.83, 1.19] | -0.06 | 0.951 |
| TST | between | 1.09 | [0.89, 1.33] | 0.79 | 0.429 |
| S-AW (Half2) (REM) | within | 0.97 | [0.87, 1.07] | -0.66 | 0.508 |
| M-AW (Half2) (REM) | within | 1.08 | [0.98, 1.19] | 1.52 | 0.128 |
| L-AW (Half2) (REM) | within | 1.16 | [1.05, 1.29] | 2.95 | **0.003\*\*** |
| S-AW (Half2) (NREM) | within | 1.05 | [0.95, 1.17] | 0.95 | 0.342 |
| M-AW (Half2) (NREM) | within | 1.02 | [0.92, 1.13] | 0.42 | 0.677 |
| L-AW (Half2) (NREM) | within | 1.06 | [0.96, 1.17] | 1.11 | 0.269 |
| TST | within | 1.22 | [1.08, 1.38] | 3.23 | **0.001\*\*** |
| PREM | other | 1.99 | [1.52, 2.60] | 5.05 | **<0.001\*\*\*** |
| SEX | other | 1.19 | [0.81, 1.75] | 0.90 | 0.369 |
| AGE | other | 1.09 | [0.89, 1.32] | 0.81 | 0.416 |
| DATASET | other | 2.23 | [1.46, 3.41] | 3.71 | **<0.001\*\*\*** |
| DAY | other | 0.80 | [0.72, 0.88] | -4.42 | **<0.001\*\*\*** |

Type: Between = between-person (person-mean) component; Within = within-person deviation from the person mean; Other = intercept and non-decomposed covariates. Continuous predictors were standardized (mean = 0, SD = 1). Sex was contrast-coded (-0.5 = male, +0.5 = female). Asterisks denote significance: \*p < .05, \*\*p < .01, \*\*\*p < .001. DATASET = Dataset 3 vs 2; DAY = study day; TST = total sleep time; S-AW = short awakening count; M-AW = medium awakening count; L-AW = long awakening count; PREM = final awakening stage (PREM); REM/NREM = sleep stage; Half1/Half2 = first/second half of the night.

**Table 27.** Model E2 (GLMM results for CD vs. WD)

| Effect (fixed) | Type | Odds ratio | 95% CI | Statistic | p-value |
| --- | --- | --- | --- | --- | --- |
| Intercept | other | 2.22 | [1.82, 2.71] | 7.83 | **<0.001\*\*\*** |
| Subj-AW | between | 1.14 | [0.95, 1.37] | 1.45 | 0.148 |
| Subj-TST | between | 1.16 | [0.97, 1.40] | 1.63 | 0.102 |
| Subj-AW | within | 1.04 | [0.94, 1.15] | 0.79 | 0.431 |
| Subj-TST | within | 1.31 | [1.18, 1.45] | 5.15 | **<0.001\*\*\*** |
| SEX | other | 1.09 | [0.75, 1.57] | 0.45 | 0.652 |
| AGE | other | 1.04 | [0.86, 1.26] | 0.40 | 0.691 |
| DATASET | other | 2.66 | [1.73, 4.08] | 4.48 | **<0.001\*\*\*** |
| DAY | other | 0.80 | [0.72, 0.88] | -4.39 | **<0.001\*\*\*** |

Type: Between = between-person (person-mean) component; Within = within-person deviation from the person mean; Other = intercept and non-decomposed covariates. Continuous predictors were standardized (mean = 0, SD = 1). Sex was contrast-coded (-0.5 = male, +0.5 = female). Asterisks denote significance: \*p < .05, \*\*p < .01, \*\*\*p < .001. DAY = study day; Subj-TST = subjective sleep duration; Subj-AW = subjective awakening count.

##### Models Summaries

**Table 28.** Model summaries for content recall.

| Model | Specification | N | ICC | Conditional R<sup>2</sup> | Marginal R<sup>2</sup> |
| --- | --- | --- | --- | --- | --- |
| M1 | Total awakening count + covariates | 2,365 | 0.13 | 0.08 | 0.06 |
| M2 | Awakening counts by duration + covariates | 2,365 | 0.14 | 0.08 | 0.06 |
| M3 | Awakening counts by duration and sleep stage | 2,365 | 0.10 | 0.10 | 0.07 |
| M4 | M3 + final REM awakening% + total sleep time | 2,365 | 0.10 | 0.12 | 0.08 |
| EM1 | M4 in second night-half | 2,365 | 0.12 | 0.12 | 0.09 |
| EM2 | Subjective sleep measures+ covariates | 2327 | 0.13 | 0.09 | 0.06 |

Incremental variance explained by mixed-effects models for CD vs. WD. Each model was controlled for sex, age, dataset, and day (since diary start).

## Supplementary Materials

### Sensitivity Analysis Results

For a sensitivity analysis applying a stricter >90% threshold, 622 participants were included (56.3% female; mean age 33.9 ± 2.8 years), yielding 1,220 wave-level observations (wave 1, n = 501; wave 2, n = 373; wave 3, n = 346). Wave-level observations were calculated as participant-level averages across all usable nights within each wave. Participants contributed 3–8 usable nights per wave (primary: mean nights per participant per wave 5.3 ± 1.4; sensitivity: 5.0 ± 1.4).

#### Main Models

**Table S1.**
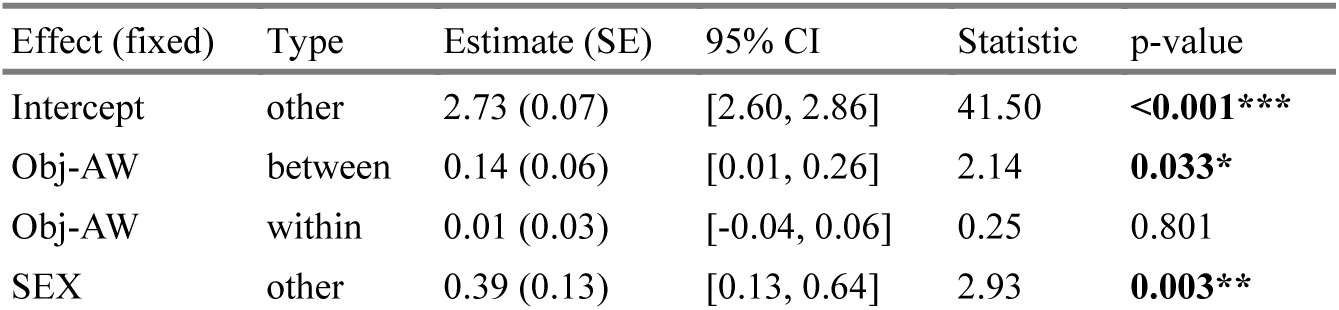
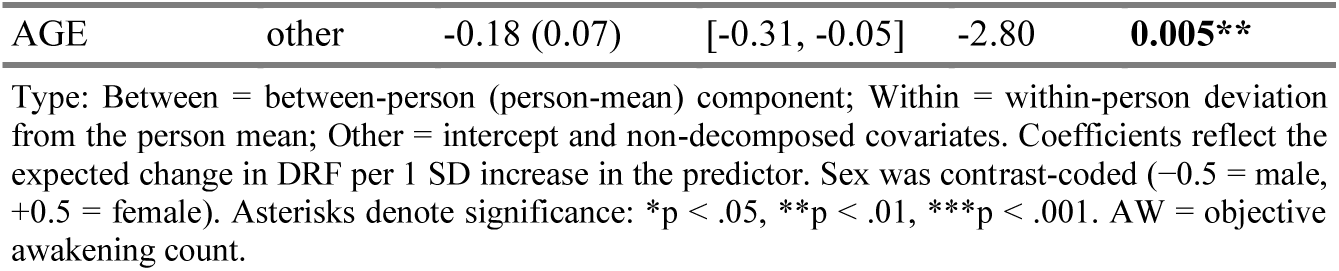
Model 1 (LME results for DRF)

| Effect (fixed) | Type | Estimate (SE) | 95% CI | Statistic | p-value |
| --- | --- | --- | --- | --- | --- |
| Intercept | other | 2.73 (0.07) | [2.60, 2.86] | 41.50 | **<0.001\*\*\*** |
| Obj-AW | between | 0.14 (0.06) | [0.01, 0.26] | 2.14 | **0.033\*** |
| Obj-AW | within | 0.01 (0.03) | [-0.04, 0.06] | 0.25 | 0.801 |
| SEX | other | 0.39 (0.13) | [0.13, 0.64] | 2.93 | **0.003\*\*** |
| AGE | other | -0.18 (0.07) | [-0.31, -0.05] | -2.80 | **0.005\*\*** |

Type: Between = between-person (person-mean) component; Within = within-person deviation from the person mean; Other = intercept and non-decomposed covariates. Coefficients reflect the expected change in DRF per 1 SD increase in the predictor. Sex was contrast-coded (-0.5 = male, +0.5 = female). Asterisks denote significance: \*p < .05, \*\*p < .01, \*\*\*p < .001. AW = objective awakening count.

**Table S2.** Model 2 (LME results for DRF)

| Effect (fixed) | Type | Estimate (SE) | 95% CI | Statistic | p-value |
| --- | --- | --- | --- | --- | --- |
| Intercept | other | 2.73 (0.07) | [2.60, 2.86] | 41.58 | **<0.001\*\*\*** |
| S-AW | between | 0.16 (0.08) | [-0.00, 0.32] | 1.94 | 0.052 |
| M-AW | between | -0.13 (0.10) | [-0.32, 0.07] | -1.28 | 0.203 |
| L-AW | between | 0.17 (0.08) | [0.02, 0.33] | 2.19 | **0.029\*** |
| S-AW | within | 0.01 (0.03) | [-0.05, 0.06] | 0.25 | 0.804 |
| M-AW | within | 0.06 (0.03) | [-0.00, 0.13] | 1.85 | 0.064 |
| L-AW | within | -0.06 (0.03) | [-0.12, -0.00] | -2.05 | **0.040\*** |
| SEX | other | 0.39 (0.13) | [0.13, 0.64] | 2.93 | **0.003\*\*** |
| AGE | other | -0.18 (0.07) | [-0.31, -0.05] | -2.78 | **0.006\*\*** |

Type: Between = between-person (person-mean) component; Within = within-person deviation from the person mean; Other = intercept and non-decomposed covariates. Coefficients reflect the expected change in DRF per 1 SD increase in the predictor. Sex was contrast-coded (-0.5 = male, +0.5 = female). Asterisks denote significance: \*p < .05, \*\*p < .01, \*\*\*p < .001. S-AW = short awakening count; M-AW = medium awakening count; L-AW = long awakening count; REM/NREM = sleep stage; Half1/Half2 = first/second half of the night.

**Table S3.** Model 3 (LME results for DRF)

| Effect (fixed) | Type | Estimate (SE) | 95% CI | Statistic | p-value |
| --- | --- | --- | --- | --- | --- |
| Intercept | other | 2.73 (0.07) | [2.60, 2.86] | 41.65 | **<0.001\*\*\*** |
| S-AW (REM) | between | -0.02 (0.09) | [-0.19, 0.15] | -0.22 | 0.826 |
| M-AW (REM) | between | 0.00 (0.10) | [-0.19, 0.20] | 0.03 | 0.976 |
| L-AW (REM) | between | 0.17 (0.09) | [-0.01, 0.35] | 1.91 | 0.057 |
| S-AW (NREM) | between | 0.19 (0.09) | [0.02, 0.36] | 2.17 | **0.030\*** |
| M-AW (NREM) | between | -0.13 (0.10) | [-0.33, 0.07] | -1.31 | 0.189 |
| L-AW (NREM) | between | 0.05 (0.09) | [-0.13, 0.22] | 0.52 | 0.602 |
| S-AW (REM) | within | -0.01 (0.03) | [-0.07, 0.05] | -0.37 | 0.715 |
| M-AW (REM) | within | 0.03 (0.03) | [-0.04, 0.09] | 0.86 | 0.392 |
| L-AW (REM) | within | -0.04 (0.03) | [-0.10, 0.02] | -1.36 | 0.173 |
| S-AW (NREM) | within | 0.02 (0.03) | [-0.04, 0.07] | 0.65 | 0.517 |
| M-AW (NREM) | within | 0.06 (0.03) | [-0.01, 0.12] | 1.70 | 0.089 |
| L-AW (NREM) | within | -0.03 (0.03) | [-0.09, 0.04] | -0.83 | 0.408 |
| SEX | other | 0.39 (0.13) | [0.13, 0.65] | 2.97 | **0.003\*\*** |
| AGE | other | -0.18 (0.07) | [-0.31, -0.06] | -2.81 | **0.005\*\*** |

Type: Between = between-person (person-mean) component; Within = within-person deviation from the person mean; Other = intercept and non-decomposed covariates. Coefficients reflect the expected change in DRF per 1 SD increase in the predictor. Sex was contrast-coded (-0.5 = male, +0.5 = female). Asterisks denote significance: \*p < .05, \*\*p < .01, \*\*\*p < .001. S-AW = short awakening count; M-AW = medium awakening count; L-AW = long awakening count; REM/NREM = sleep stage; Half1/Half2 = first/second half of the night.

**Table S4.** Model 4 (LME results for DRF)

| Effect (fixed) | Type | Estimate (SE) | 95% CI | Statistic | p-value |
| --- | --- | --- | --- | --- | --- |
| Intercept | other | 2.73 (0.07) | [2.60, 2.86] | 41.70 | **<0.001\*\*\*** |
| S-AW (REM) | between | -0.06 (0.09) | [-0.24, 0.12] | -0.69 | 0.492 |
| M-AW (REM) | between | -0.01 (0.10) | [-0.21, 0.18] | -0.13 | 0.898 |
| L-AW (REM) | between | 0.20 (0.09) | [0.02, 0.38] | 2.16 | **0.032\*** |
| S-AW (NREM) | between | 0.16 (0.09) | [-0.01, 0.34] | 1.81 | 0.070 |
| M-AW (NREM) | between | -0.13 (0.10) | [-0.33, 0.07] | -1.23 | 0.219 |
| L-AW (NREM) | between | 0.08 (0.09) | [-0.10, 0.26] | 0.90 | 0.368 |
| TST | between | 0.12 (0.08) | [-0.04, 0.27] | 1.50 | 0.135 |
| FREM% | between | 0.06 (0.07) | [-0.08, 0.20] | 0.79 | 0.429 |
| S-AW (REM) | within | -0.02 (0.03) | [-0.08, 0.04] | -0.79 | 0.429 |
| M-AW (REM) | within | 0.02 (0.03) | [-0.04, 0.09] | 0.72 | 0.474 |
| L-AW (REM) | within | -0.04 (0.03) | [-0.10, 0.03] | -1.12 | 0.262 |
| S-AW (NREM) | within | 0.01 (0.03) | [-0.05, 0.06] | 0.27 | 0.790 |
| M-AW (NREM) | within | 0.06 (0.03) | [-0.01, 0.12] | 1.76 | 0.079 |
| L-AW (NREM) | within | -0.01 (0.03) | [-0.07, 0.06] | -0.19 | 0.846 |
| TST | within | 0.05 (0.03) | [-0.01, 0.11] | 1.55 | 0.121 |
| FREM% | within | -0.00 (0.03) | [-0.05, 0.05] | -0.01 | 0.995 |
| SEX | other | 0.39 (0.13) | [0.13, 0.66] | 2.94 | **0.003\*\*** |
| AGE | other | -0.19 (0.07) | [-0.32, -0.06] | -2.93 | **0.004\*\*** |

Type: Between = between-person (person-mean) component; Within = within-person deviation from the person mean; Other = intercept and non-decomposed covariates. Coefficients reflect the expected change in DRF per 1 SD increase in the predictor. Sex was contrast-coded (-0.5 = male, +0.5 = female). Asterisks denote significance: \*p < .05, \*\*p < .01, \*\*\*p < .001. TST = total sleep time; S-AW = short awakening count; M-AW = medium awakening count; L-AW = long awakening count; FREM% = percentage of final awakenings preceded by REM; REM/NREM = sleep stage; Half1/Half2 = first/second half of the night.

#### Exploratory Models

**Table S5.**
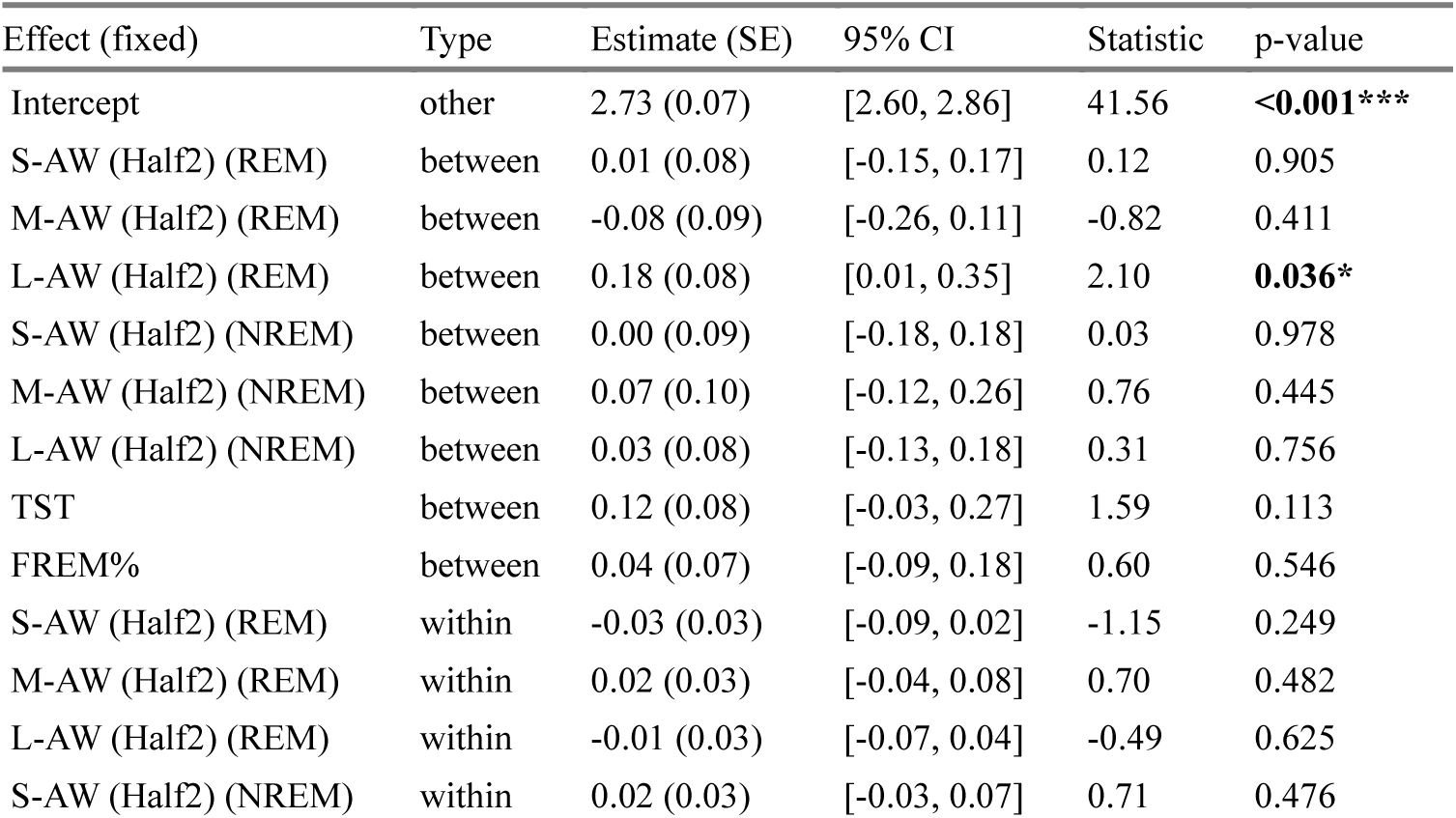
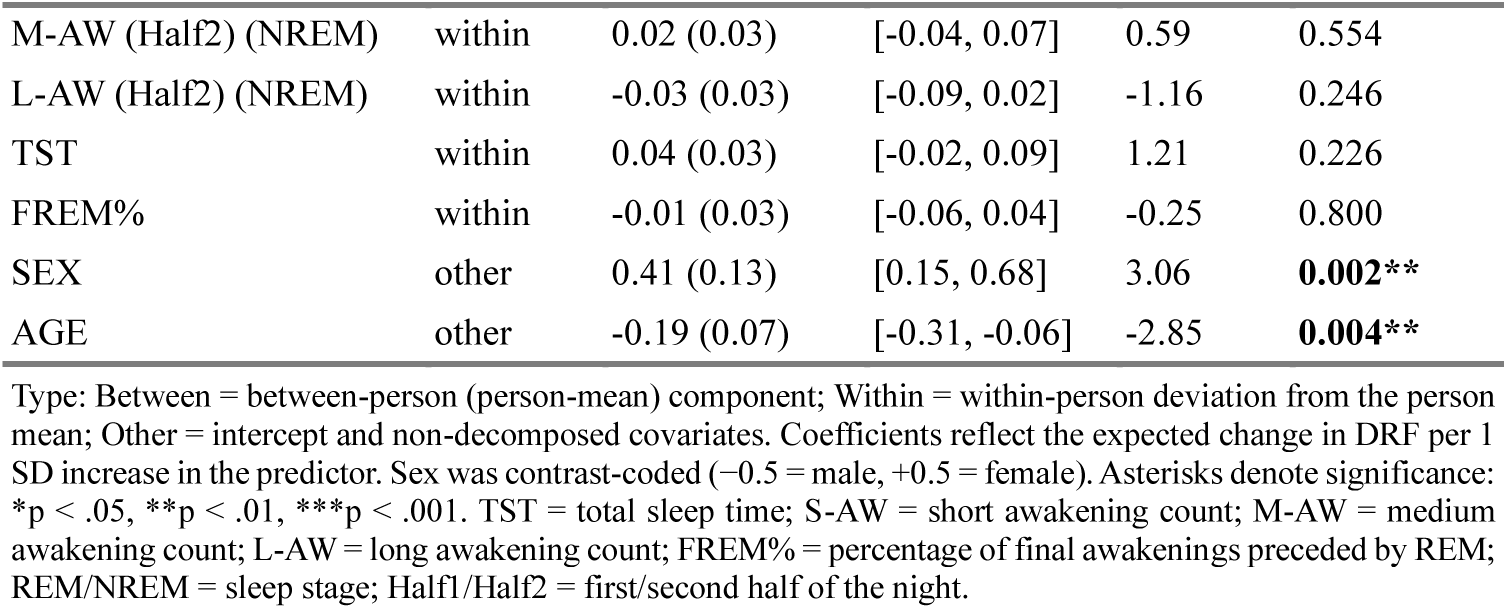
Model E1 (LME results for DRF)

| Effect (fixed) | Type | Estimate (SE) | 95% CI | Statistic | p-value |
| --- | --- | --- | --- | --- | --- |
| Intercept | other | 2.73 (0.07) | [2.60, 2.86] | 41.56 | **<0.001\*\*\*** |
| S-AW (Half2) (REM) | between | 0.01 (0.08) | [-0.15, 0.17] | 0.12 | 0.905 |
| M-AW (Half2) (REM) | between | -0.08 (0.09) | [-0.26, 0.11] | -0.82 | 0.411 |
| L-AW (Half2) (REM) | between | 0.18 (0.08) | [0.01, 0.35] | 2.10 | **0.036\*** |
| S-AW (Half2) (NREM) | between | 0.00 (0.09) | [-0.18, 0.18] | 0.03 | 0.978 |
| M-AW (Half2) (NREM) | between | 0.07 (0.10) | [-0.12, 0.26] | 0.76 | 0.445 |
| L-AW (Half2) (NREM) | between | 0.03 (0.08) | [-0.13, 0.18] | 0.31 | 0.756 |
| TST | between | 0.12 (0.08) | [-0.03, 0.27] | 1.59 | 0.113 |
| FREM% | between | 0.04 (0.07) | [-0.09, 0.18] | 0.60 | 0.546 |
| S-AW (Half2) (REM) | within | -0.03 (0.03) | [-0.09, 0.02] | -1.15 | 0.249 |
| M-AW (Half2) (REM) | within | 0.02 (0.03) | [-0.04, 0.08] | 0.70 | 0.482 |
| L-AW (Half2) (REM) | within | -0.01 (0.03) | [-0.07, 0.04] | -0.49 | 0.625 |
| S-AW (Half2) (NREM) | within | 0.02 (0.03) | [-0.03, 0.07] | 0.71 | 0.476 |
| M-AW (Half2) (NREM) | within | 0.02 (0.03) | [-0.04, 0.07] | 0.59 | 0.554 |
| L-AW (Half2) (NREM) | within | -0.03 (0.03) | [-0.09, 0.02] | -1.16 | 0.246 |
| TST | within | 0.04 (0.03) | [-0.02, 0.09] | 1.21 | 0.226 |
| FREM% | within | -0.01 (0.03) | [-0.06, 0.04] | -0.25 | 0.800 |
| SEX | other | 0.41 (0.13) | [0.15, 0.68] | 3.06 | **0.002\*\*** |
| AGE | other | -0.19 (0.07) | [-0.31, -0.06] | -2.85 | **0.004\*\*** |

Type: Between = between-person (person-mean) component; Within = within-person deviation from the person mean; Other = intercept and non-decomposed covariates. Coefficients reflect the expected change in DRF per 1 SD increase in the predictor. Sex was contrast-coded (-0.5 = male, +0.5 = female). Asterisks denote significance: \*p < .05, \*\*p < .01, \*\*\*p < .001. TST = total sleep time; S-AW = short awakening count; M-AW = medium awakening count; L-AW = long awakening count; FREM% = percentage of final awakenings preceded by REM; REM/NREM = sleep stage; Half1/Half2 = first/second half of the night.

**Table S6.** Model E2 (LME results for DRF)

| Effect (fixed) | Type | Estimate (SE) | 95% CI | Statistic | p-value |
| --- | --- | --- | --- | --- | --- |
| Intercept | other | 2.72 (0.07) | [2.59, 2.85] | 41.51 | **<0.001\*\*\*** |
| Subj-AW | between | 0.29 (0.07) | [0.15, 0.42] | 4.24 | **<0.001\*\*\*** |
| Subj-TST | between | 0.15 (0.07) | [0.02, 0.28] | 2.20 | **0.028\*** |
| Subj-AW | within | 0.01 (0.03) | [-0.04, 0.06] | 0.29 | 0.770 |
| Subj-TST | within | 0.03 (0.03) | [-0.02, 0.08] | 1.04 | 0.301 |
| SEX | other | 0.37 (0.13) | [0.11, 0.63] | 2.80 | **0.005\*\*** |
| AGE | other | -0.21 (0.06) | [-0.34, -0.09] | -3.33 | **<0.001\*\*\*** |

Type: Between = between-person (person-mean) component; Within = within-person deviation from the person mean; Other = intercept and non-decomposed covariates. Coefficients reflect the expected change in DRF per 1 SD increase in the predictor. Sex was contrast-coded (-0.5 = male, +0.5 = female). Asterisks denote significance: \*p < .05, \*\*p < .01, \*\*\*p < .001. Abbreviations: Subj-TST = subjective sleep duration; Subj-AW = subjective awakening count.

**Table S7.** Model E3 (LME results for DRF)

| Effect (fixed) | Type | Estimate (SE) | 95% CI | Statistic | p-value |
| --- | --- | --- | --- | --- | --- |
| Intercept | other | 2.68 (0.08) | [2.52, 2.84] | 33.05 | **<0.001\*\*\*** |
| Subj-AW | between | 0.26 (0.09) | [0.09, 0.44] | 2.92 | **0.004\*\*** |
| Subj-TST | between | -0.00 (0.09) | [-0.17, 0.17] | -0.01 | 0.991 |
| ALCO | between | 0.11 (0.09) | [-0.07, 0.28] | 1.22 | 0.224 |
| COFF | between | -0.13 (0.09) | [-0.31, 0.04] | -1.49 | 0.137 |
| CANN | between | 0.16 (0.07) | [0.02, 0.30] | 2.21 | **0.027\*** |
| DEP | between | -0.06 (0.12) | [-0.28, 0.17] | -0.49 | 0.627 |
| Anx-State | between | 0.08 (0.12) | [-0.15, 0.30] | 0.65 | 0.513 |
| AM-L | between | 0.06 (0.09) | [-0.11, 0.23] | 0.72 | 0.470 |
| AM-S | between | 0.03 (0.09) | [-0.15, 0.21] | 0.37 | 0.712 |
| Subj-AW | within | 0.03 (0.03) | [-0.04, 0.09] | 0.87 | 0.387 |
| Subj-TST | within | 0.03 (0.03) | [-0.03, 0.09] | 1.06 | 0.291 |
| ALCO | within | 0.06 (0.03) | [0.00, 0.13] | 1.97 | **0.049\*** |
| COFF | within | 0.01 (0.03) | [-0.05, 0.07] | 0.24 | 0.812 |
| CANN | within | -0.01 (0.06) | [-0.12, 0.10] | -0.23 | 0.817 |
| DEP | within | -0.05 (0.03) | [-0.11, 0.02] | -1.36 | 0.175 |
| Anx-State | within | 0.00 (0.03) | [-0.06, 0.07] | 0.05 | 0.963 |
| AM-L | within | -0.02 (0.03) | [-0.08, 0.05] | -0.53 | 0.596 |
| AM-S | within | -0.03 (0.03) | [-0.09, 0.03] | -1.09 | 0.276 |
| VWM | other | 0.08 (0.08) | [-0.09, 0.24] | 0.94 | 0.347 |
| AWM | other | 0.08 (0.08) | [-0.08, 0.25] | 1.02 | 0.308 |
| SEX | other | 0.22 (0.17) | [-0.11, 0.55] | 1.30 | 0.193 |
| AGE | other | -0.20 (0.08) | [-0.36, -0.04] | -2.44 | **0.015\*** |

Type: Between = between-person (person-mean) component; Within = within-person deviation from the person mean; Other = intercept and non-decomposed covariates. Coefficients reflect the expected change in DRF per 1 SD increase in the predictor. Sex was contrast-coded (-0.5 = male, +0.5 = female). Asterisks denote significance: \* $p < .05$ , \*\* $p < .01$ , \*\*\* $p < .001$ . Abbreviations: Subj-TST = subjective sleep duration; Subj-AW = subjective awakening count; COFF = proportion of beeps with coffee intake; ALCO = proportion of beeps with alcohol intake; CANN = proportion of beeps with cannabis intake; Anx-State = state anxiety score; DEP = depression score; AM-L = associative memory (long) accuracy; AM-S = associative memory (short) accuracy; VWM = visual working memory (forward); AWM = auditory working memory (forward).

### Exclusion flow

**Table S8.** Dataset 1 (75%) exclusion flow.

| Step | Description | Participants excluded | Participants remaining | Nights excluded | Nights remaining |
| --- | --- | --- | --- | --- | --- |
| 0 | TST available (entry set) | — | 817 | — | 12,240 |
| 1 | Exclude TST < 180 min | 16 | 801 | 2,611 | 9,629 |
| 2 | Exclude scorable% < 75 or missing | 4 | 797 | 854 | 8,775 |
| 3 | Exclude final-awakening stage not REM/NREM | 0 | 797 | 169 | 8,606 |
| 4 | Exclude Participant×Time cells with <3 nights | 82 | 715 | 553 | 8,053 |
| 5 | Match with Survey (DRF present) | 7 | 708 | 223 | 7,830 |

**Table S9.** Dataset 1 (90%) exclusion flow.

| Step | Description | Participants excluded | Participants remaining | Nights excluded | Nights remaining |
| --- | --- | --- | --- | --- | --- |
| 0 | TST available (entry set) | — | 817 | — | 12,240 |
| 1 | Exclude TST < 180 min | 16 | 801 | 2,611 | 9,629 |
| 2 | Exclude scorable% < 75 or missing | 28 | 773 | 2,507 | 7,122 |
| 3 | Exclude final-awakening stage not REM/NREM | 0 | 773 | 64 | 7,058 |
| 4 | Exclude Participant×Time cells with <3 nights | 143 | 630 | 760 | 6,298 |
| 5 | Match with Survey (DRF present) | 8 | 622 | 178 | 6,120 |

**Table S10.** Dataset 2 exclusion flow.

| Step | Description | Participants excluded | Participants remaining | Nights excluded | Nights remaining |
| --- | --- | --- | --- | --- | --- |
| 0 | Participants with available sleep data | — | 36 | — | 1,254 |
| 1 | Exclude: diaries filled after next sleep session | 0 | 36 | 51 | 1,203 |
| 2 | Exclude: nights with Fitbit end after or within 30 min before diary wakeup | 0 | 36 | 75 | 1,128 |
| 3 | Exclude: participants with <3 valid nights per Time | 0 | 36 | 0 | 1,128 |
| 4 | Exclude: participants with DRF <4 | 8 | 28 | 253 | 875 |

**Table S11.** Dataset 3 exclusion flow.

| Step | Description | Participants excluded | Participants remaining | Nights excluded | Nights remaining |
| --- | --- | --- | --- | --- | --- |
| 0 | Participants with available sleep data | — | 100 | — | 2,338 |
| 1 | Exclude: diaries filled after next sleep session | 0 | 100 | 129 | 2,209 |
| 2 | Exclude: nights with Fitbit end after or within 30 min before diary wakeup | 0 | 100 | 96 | 2,113 |
| 3 | Exclude: participants with <3 valid nights per Time | 1 | 99 | 2 | 2,111 |
| 4 | Exclude: participants with DRF <4 | 3 | 96 | 90 | 2,021 |

---

**Fig. S1:**
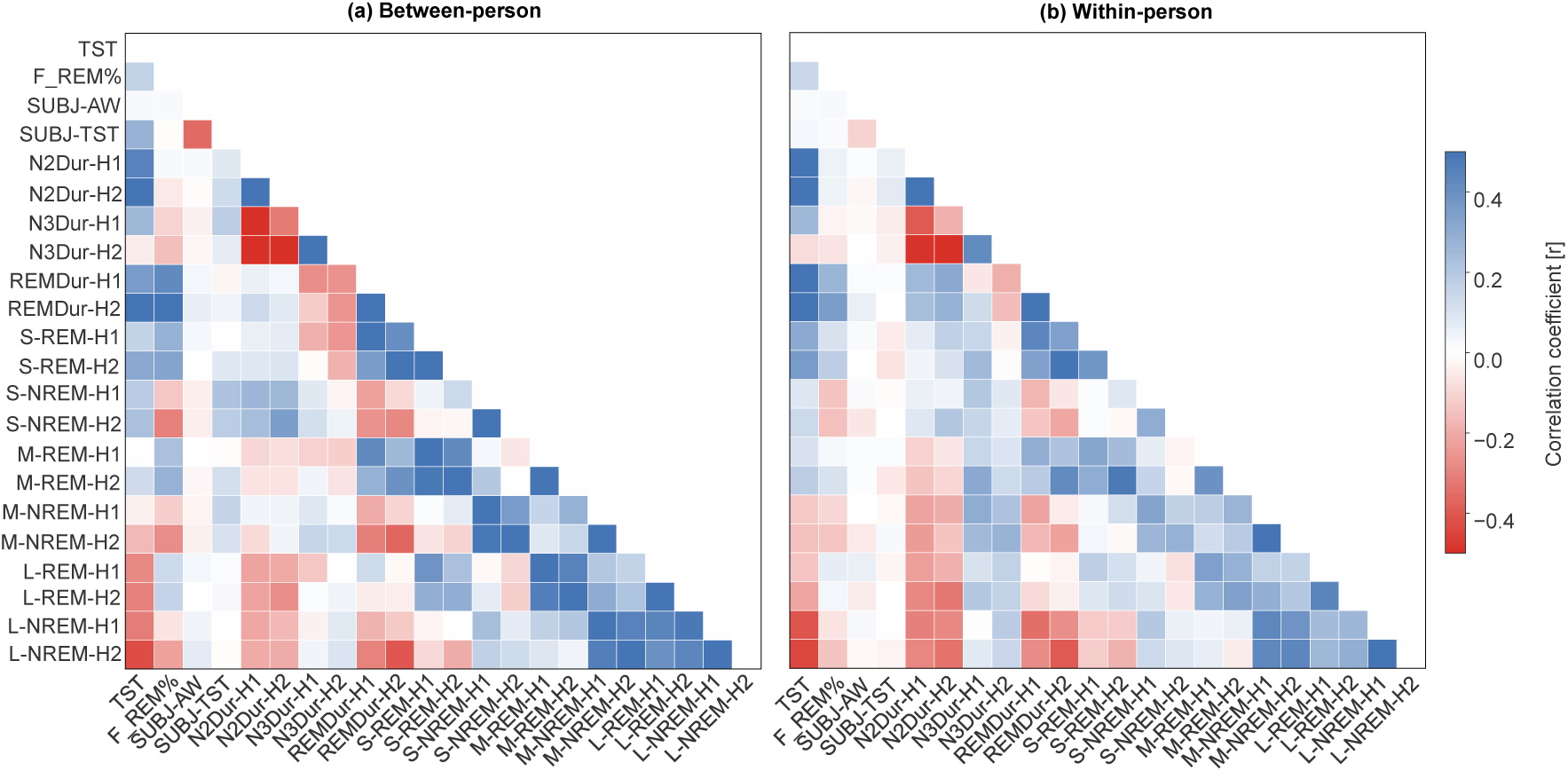
Correlations (pairwise Pearson’s r) among objective and subjective awakening and sleep measures at both within- and between-person levels in Dataset 1 Between-person (a) and within-person (b) Pearson correlation heatmaps for sleep measures. Blue indicates positive correlations and red indicates negative correlations. TST = total sleep time; F_REM% = preceding final REM percentage; SUBJ-AW = subjective number of awakenings; SUBJ-TST = subjective sleep duration; N2Dur-H1/H2 = N2 duration in the first/second half of the night; N3Dur-H1/H2 = N3 duration in the first/second half of the night; REMDur-H1/H2 = REM duration in the first/second half of the night. S, M, and L denote short, medium, and long awakenings, respectively; REM and NREM indicate sleep stage; and H1/H2 denote the first and second half of the night.

**Fig. S2:**
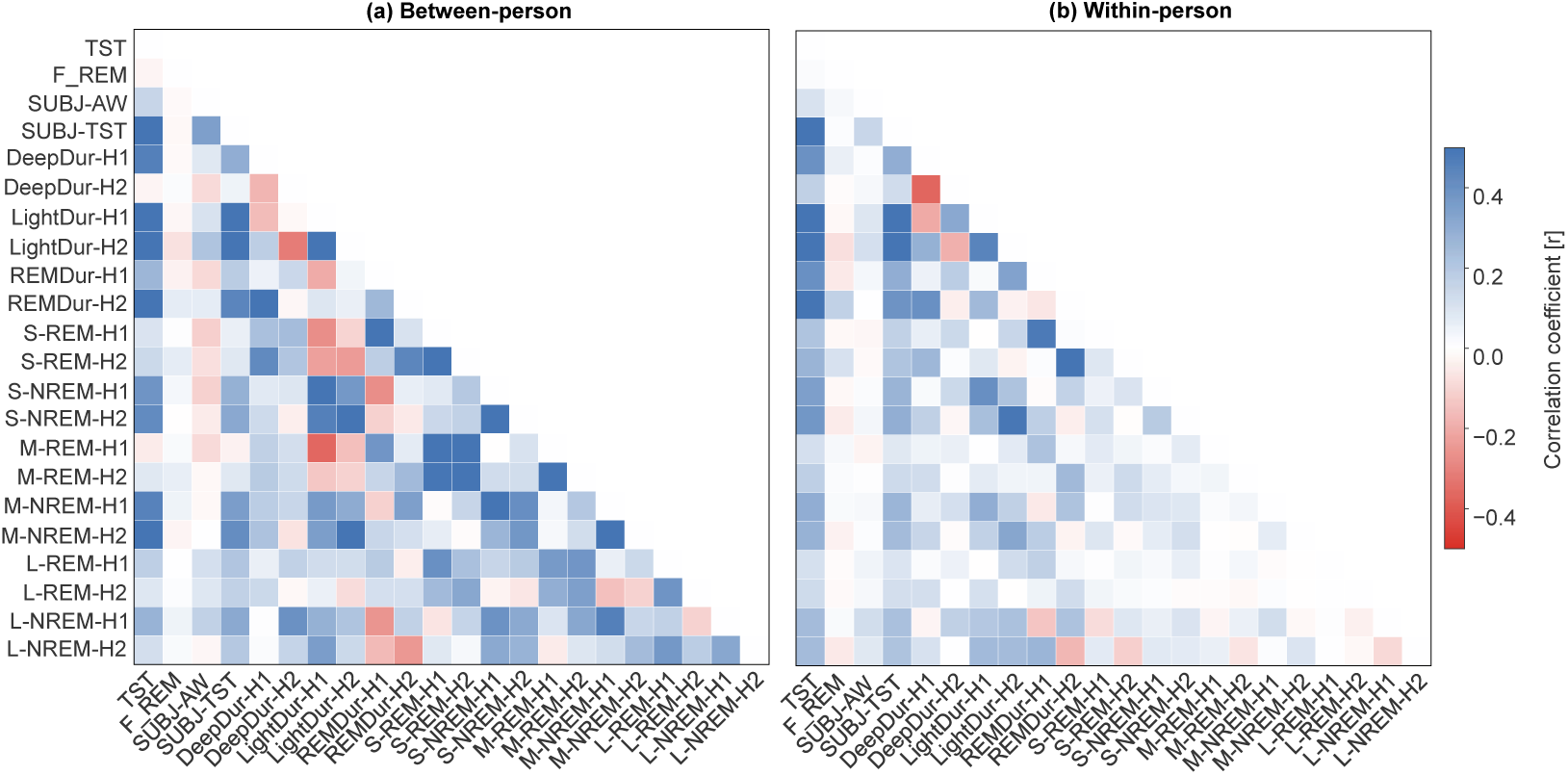
Correlations (pairwise Pearson’s r) among objective and subjective awakening and sleep measures at both within- and between-person levels in Dataset 2+3 Between-person (a) and within-person (b) Pearson correlation heatmaps for sleep variables. Blue indicates positive correlations and red indicates negative correlations. TST = total sleep time; F_REM = final awakening stage (REM vs. NREM); SUBJ-AW = subjective number of awakenings; SUBJ-TST = subjective sleep duration; DeepDur-H1/H2 = deep sleep duration in the first/second half of the night; LightDur-H1/H2 = light sleep duration in the first/second half of the night; REMDur-H1/H2 = REM sleep duration in the first/second half of the night. S, M, and L indicate short, medium, and long awakenings; REM/NREM indicate sleep stage; and H1/H2 indicate the first and second half of the night.

## Notes

### Competing Interest Statement

The authors have declared no competing interest.

